# Antioxidant supplementation blunts the proteome response to three weeks of sprint interval training preferentially in human type 2 muscle fibres

**DOI:** 10.1101/2025.01.27.634979

**Authors:** Victoria L Wyckelsma, Marta Murgia, Sigitas Kamandulis, Stefano Gastaldello, Marius Brazaitis, Audrius Snieckus, Nerijus Eimantas, Mati Pääsuke, Sebastian Edman, William Apro, Daniel C Andersson, Håkan Westerblad, Tomas Venckunas

## Abstract

Sprint interval training (SIT) is a time-efficient type of endurance training that involves large type 2 muscle fibre recruitment. Effiective antioxidant supplementation may mitigate positive training adaptations by limiting the oxidant challenge. Our aim was to test whether SIT affiects type 2 more than type 1 muscle fibres, and whether the training response is mitigated by antioxidant treatment. Young men performed three SIT sessions (6 × 30 s all-out cycling) per week for three weeks while treated with antioxidants (vitamin C, 1 g/day; vitamin E, 235 mg/day) or placebo. Vastus lateralis biopsies were taken to measure (i) activation of genes for reactive oxygen/nitrogen species (ROS) sensors and inflammatory mediators with quantitative RT PCR and (ii) fibre type-specific proteome adaptations using mass spectrometry-based proteomics. Vitamin treatment decreased the upregulation of genes for ROS sensors and inflammatory regulators during the first SIT session. The three weeks of SIT caused generally larger proteome adaptations in type 2 than in type 1 fibres, and this included larger increases in abundance of proteins involved in mitochondrial energy production. Vitamin treatment blunted the SIT-induced proteome adaptations, whereas it did not affiect the training-induced improvement in maximal cycling performance. In conclusion, (i) the large type 2 fibre recruitment and resulting proteome adaptations are instrumental to the effiectiveness of SIT, and (ii) antioxidant supplementation counteracts positive muscular adaptations to SIT, which would blunt any improvement in submaximal endurance performance, whereas it does not affiect the improvement in maximal cycling performance, where O2 delivery to muscle would be limiting.

## INTRODUCTION

A general concept of endurance training is to stress energy metabolism and thereby trigger adaptations towards increased aerobic capacity and improved performance during endurance-type exercises. There are many ways to perform endurance training, ranging from high-volume exercise at very low intensity to low-volume sprint interval training at very high intensity. Nevertheless, it is well-established that optimal endurance performance requires a combination of low-and high-intensity exercises (Sperlich *et al*., 2023).

From a skeletal muscle fibre perspective, prolonged low-intensity and short-lasting high-intensity endurance exercises present markedly diffierent challenges. Mainly slow-twitch type 1 fibres are recruited during prolonged low-intensity exercise, whereas both type 1 fibres and fast-twitch type 2 fibres are recruited during high-intensity interval training (Vøllestad & Blom, 1985). Accordingly, type 2 fibres would be expected to be more affiected by high-than low-intensity training (Henriksson & Reitman, 1976; Dudley *et al*., 1982; MacInnis & Gibala, 2017; Reisman *et al*., 2024b). Diffierences in fibre type recruitment complicate the interpretation of human data of exercise-induced changes in energy metabolites as well as in gene and protein expression, including various omics, since these are mostly obtained at the whole muscle level (MacInnis & Gibala, 2017; Wyckelsma *et al*., 2017; Reisman *et al*., 2024b).

An oxidative dominance in the relation between reactive oxygen/nitrogen species (ROS) and their antioxidant defence systems was classically termed “oxidative stress” and linked to oxidative damage to cells and organs (Sies & Cadenas, 1985). This concept has since been qualified and it is now well-established that while excessive oxidant challenges are deleterious, physiological challenges have important signalling roles and are essential for normal cell function (Sies *et al*., 2017). For instance, bursts of increased ROS production during endurance exercise can trigger beneficial skeletal muscle adaptations (Margaritelis *et al*., 2018; Jordan *et al*., 2021; Li *et al*., 2022; Powers *et al*., 2024). Accordingly, many studies show mitigated endurance training responses in skeletal muscle with antioxidant supplementation (Gomez-Cabrera *et al*., 2008; Ristow *et al*., 2009; Strobel *et al*., 2011; Paulsen *et al*., 2014), although there are also studies showing no blunting or even positive effiect of antioxidant treatment (Aguilo *et al*., 2007; Yfanti *et al*., 2010; Flockhart *et al*., 2023).

A shift in the redox balance towards oxidation is implicated in the dual effiects of inflammatory signalling in skeletal muscle: long-lasting systemic increases of inflammatory mediators, such as various cytokines, are associated with skeletal muscle dysfunctions and atrophy, whereas transient exercise-induced muscular increases of the same cytokines may contribute to positive training responses (Munoz-Canoves *et al*., 2013; Raschke & Eckel, 2013; Peake *et al*., 2015). Moreover, increased ROS has been shown to affiect the sarcoplasmic reticulum (SR) Ca^2+^ release channel complex centred at the tetrameric ryanodine receptor 1 (RyR1) protein, for instance, by causing dissociation of the channel stabilizing FK506-binding protein 12 (FKBP12) or fragmentation of the RyR1 (Bellinger *et al*., 2008a; Place *et al*., 2015). Major ROS-mediated changes in the RyR1 protein complex, resulting in severe SR Ca^2+^ leak, have been linked to muscle weakness in overtraining (Aydin *et al*., 2008; Bellinger *et al*., 2008b), whereas limited SR Ca^2+^ leak can stimulate mitochondrial biogenesis and thereby increase muscular endurance (Ojuka *et al*., 2002; Aydin *et al*., 2008; Ivarsson *et al*., 2019).

In the present study, young men performed three weeks of SIT (30 s all-out cycling bouts, three sessions per week) while consuming high-dose antioxidant supplement (Vitamin C and E) or placebo. We hypothesised that SIT triggers generally larger changes in protein expression in type 2 than in type 1 fibres. Moreover, we hypothesised that the antioxidant supplementation mitigates SIT-induced activations of ROS-sensitive genes, thereby limiting beneficial muscle fibre adaptations towards increased antioxidant capacity, improved SR Ca^2+^-handling, and enhanced oxidative capacity. To test these hypotheses, we used (i) quantitative real-time PCR to measure changes in gene expression induced during the first and last SIT sessions and (ii) fibre type-specific mass spectrometry-based proteomics to assess adaptations during the three weeks of SIT. In addition, we tested whether SIT-induced diffierences in muscle protein expression between vitamin and placebo treatments affiected physiological parameters, i.e. power output and maximal oxygen uptake (VO2max) during exhaustive cycling exercise and forces during the subsequent recovery.

## METHODS

### Ethical approval

All participants provided written informed consent. The experimental procedures were conducted in accordance with the most recent version of the Declaration of Helsinki. The study was approved by the Kaunas Regional Biomedical Research Ethics Committee (BE-2-35) and Research Ethics Committee of the University of Tartu (267/T-9, 2017).

### Participants and study outline

Recreationally active young men participated in this double-blind study; none of the participants were engaged in any structured sport training program or taking any medications known to affect physical function. Participants were randomly divided into either a vitamin (n = 11) or a placebo (n = 11) group; their characteristics at the start of the study are presented in Table 1. Vitamins were provided in the form of oral vitamin C (1 g daily) and vitamin E (235 mg daily) (Paulsen *et al*., 2014; Wyckelsma *et al*., 2020). Treatments were initiated 7 days before the first SIT session. Vitamin C (or placebo) was taken as 500 mg in the morning and 500 mg in the evening, while vitamin E (or placebo) was taken as a single dose either with the morning or the evening meal. On the training days, tablets were taken at least one hour before the training session.

**Table 1.** Characteristics of participants at the start of the study.

|  | **Vitamin C + E (n = 11)** | **Placebo (n = 11)** |
| --- | --- | --- |
| Age (years) | 23.5 ± 3.6 | 26.4 ± 5.4 |
| Height (cm) | 183.5 ± 8.3 | 179.7 ± 9.0 |
| Body mass (kg) | 86.2 ± 14.6 | 80.4 ± 13.1 |
| BMI (kg m<sup>-2</sup>) | 25.6 ± 4.0 | 25.0 ± 2.5 |
| *Incremental cycling test* |  |  |
| VO<sub>2max</sub> (ml kg<sup>-1</sup> min<sup>-1</sup>) | 41.9 ± 6.5 | 42.5 ± 7.6 |
| Maximal power (W kg<sup>-1</sup>) | 3.74 ± 0.66 | 3.74 ± 0.53 |
| Maximal heart rate (min<sup>-1</sup>) | 191.4 ± 6.1 | 187.9 ± 8.4 |

Values are mean ± SD. BMI, body mass index; VO<sub>2max</sub>, maximal oxygen uptake.

Training consisted of nine sessions (3 sessions/week for 3 weeks). Each SIT session started with warm-up consisting of 8 min cycling at a power (W) equal to the individual’s body mass (kg). Then followed 4-6 repetitions of 30-s all-out cycling bouts (Wingate tests) with 4 min of rest between bouts. Sessions 1 and 7–9 were composed of 6 sprints; sessions 2-3 were composed of 4 sprints, and sessions 4–6 were composed of 5 sprints (Schlittler *et al*., 2019; Wyckelsma *et al*., 2020). A cycle ergometer with continuous power recording was used to quantify the amount of work produced during each SIT session in six of eleven participants in each group. Blood lactate was measured in fingertip blood samples taken 5 min after the first and last SIT sessions using a portable lactate analysing unit (ProTM LT-1730, Arkray Inc., Kyoto, Japan). Muscle function testing was performed before and directly (∼2 min), 1 hour, and 24 hours after the first and last SIT sessions. Vastus lateralis muscle biopsies were excised before, 1 hour and 24 hours after the first and last SIT session.

Participants were told to maintain their regular diet and no food intake for at least two hours before the first and the last SIT sessions as well as before the pre-and post-training testing sessions. Participants were familiarized with the SIT training and experimental procedures for muscle function testing on a separate occasion before the actual testing.

Participants visited the laboratory 7 days prior to the first SIT session and 48 h after the last SIT session for assessment of maximal power output and VO2max using a standard incremental test to exhaustion on a cycle ergometer (Ergometrics 800S, ErgoLine, Medical Measurement Systems, Binz, Germany). Expired gases were measured breath-by-breath with a calibrated mobile testing system (Oxycon Mobile, Jaeger/VIASYS Healthcare, Hoechberg, Germany), and heart rate (HR) was measured with an HR monitor (S625X, Polar Electro, Kempele, Finland). The cycle ergometer test started with 3 min of cycling at 40 W after which load was increased by 5 W every 10 s. Participants were told to pedal at a cadence of 70 rpm and the test was continued until they could no longer pedal at this cadence. VO2max and maximal HR were calculated as the highest average over 20 consecutive seconds.

### Muscle function

Maximal voluntary contraction (MVC) and electrically stimulated isometric force of the dominant leg knee extensors (m. quadriceps) were measured using an isokinetic dynamometer (System 3, Biodex Medical Systems, Shirley, USA). After warm-up consisting of 8 min of cycling at 80 W followed by 10 easy squats, participants were seated in the dynamometer chair with the knee joint adjusted at an angle of 70° (full knee extension = 0°). To minimize the changes in body position, shank, trunk, and shoulders were tightly fastened by belts with Velcro straps. During MVC measurements, participants were verbally encouraged to exert maximal force and maintain this force for ∼3 s. They received immediate feedback from a screen showing the force production curve. MVC was determined as the largest force of two attempts with 1 min rest between contractions.

Transcutaneous muscle stimulation was applied via a pair of carbonized rubber electrodes covered on the inner surface with a thin layer of electrode gel (ECG–EEG Gel, Medigel, Modi’in, Israel). One electrode (6 cm × 11 cm) was placed transversely across the width of the proximal segment of the m. quadriceps close to the inguinal ligament and the other electrode (6 cm × 20 cm) was placed above the distal segment of the m. quadriceps close to the patella. An electrical stimulator (MG 440, Medicor, Budapest, Hungary) was used to deliver 1 ms square-wave supramaximal pulses (voltage 10% higher of that pre-determined to elicit peak torque to a single pulse). Peak torque in response to a 1 s train of pulses given at 20 Hz was measured. Change in the 20 Hz/MVC torque ratio was used to evaluate the extent of prolonged low-frequency force depression (PLFFD)(Allen *et al*., 2008).

### Muscle biopsies

We used previously described and validated procedures for taking needle muscle biopsies (Magistris *et al*., 1998). Briefly, after skin sterilization and local anesthesia, a 1–2-mm-long skin cut was made with the tip of a scalpel mid-way over the vastus lateralis muscles of the non-dominant leg using an automatic biopsy device (Bard Biopsy Instrument, Bard Radiology, Covington, GA, USA). A 14-gauge disposable needle was inserted through the cut until the fascia was pierced and the needle was advanced ∼2 cm into the muscle, perpendicular to the muscle fibres. Two to three samples (∼15 mg each) were collected from one puncture site at each time point; different puncture sites were used for biopsies taken before, and 1 and 24 hours after the first and last SIT sessions. A local compression was then applied on the biopsy site for a few minutes to prevent hematoma. Muscle samples were immediately frozen in liquid nitrogen and stored at-80°C. Due to limited muscle tissue availability caused by a freezer breakdown, we prioritized measurements of gene expression in biopsies taken before and 1 hour after the first and last SIT sessions, whereas protein expression was measured in biopsies taken before the first and 24 hours after last SIT sessions. This prioritization was based on the fact that after exercise, changes in mRNA levels occur more rapidly than changes in protein content (Perry *et al*., 2010), and we have previously observed decreased mRNA levels of exercise-responsive genes 24 hours after a SIT session (Place *et al*., 2015).

### Gene expression analysis

Total RNA was isolated from muscle biopsies using TRIzol Reagent solution (15596026, Thermo Fisher Scientific, Carlsbad, CA, USA) following the manufacturer instructions. Genomic DNA contaminations from RNA preparations were removed by incubation with DNase I, RNAse free (EN0521, Thermo Fisher Scientific, Carlsbad, CA, USA) for 1h at 37 °C. The correspondent cDNAs were produced using both oligo (dT)18 and random hexamer primers by following the instruction of RevertAID H Minus First strand cDNA synthesis Kit (K1632, Thermo Fisher Scientific, Carlsbad, CA, USA). Quantitative real-time PCR reactions were performed with 100 ng of cDNA template using SYBR Green Master Mix (A25741, Life Technologies, Carlsbad, CA, USA) in a final volume of 20 μl. The essay was performed with QuantStudio 3 Real-Time PCR Systems, using the following cycling program: initial 50 °C 2 min, denaturation 95°C 10 min, followed by 40 cycles of 95°C for 15 seconds and 60°C for 1 min. A final step of melting curve between 65°C to 90°C, 1°C/sec temperature speed was incorporated. Table 2 shows the primers used with a stock concentration of 10 µM. Hypoxanthine-guanine phosphoribosyltransferase (HPRT1) was used as the housekeeping control gene, and it did not change with any intervention. Fold change relative to the gene expression was calculated as 2-ΔΔCt where: ΔΔCt = ΔCt (target gene 1 hour after SIT session) - ΔCt (target gene before SIT session). ΔCt = Ct[Target] - Ct[Housekeeping], according to the Minimum Information for Publication of Quantitative Real-Time PCR Experiments (MIQE) guideline. All single samples were analysed in triplicate and the mean value was used in subsequent analyses.

**Table 2.**
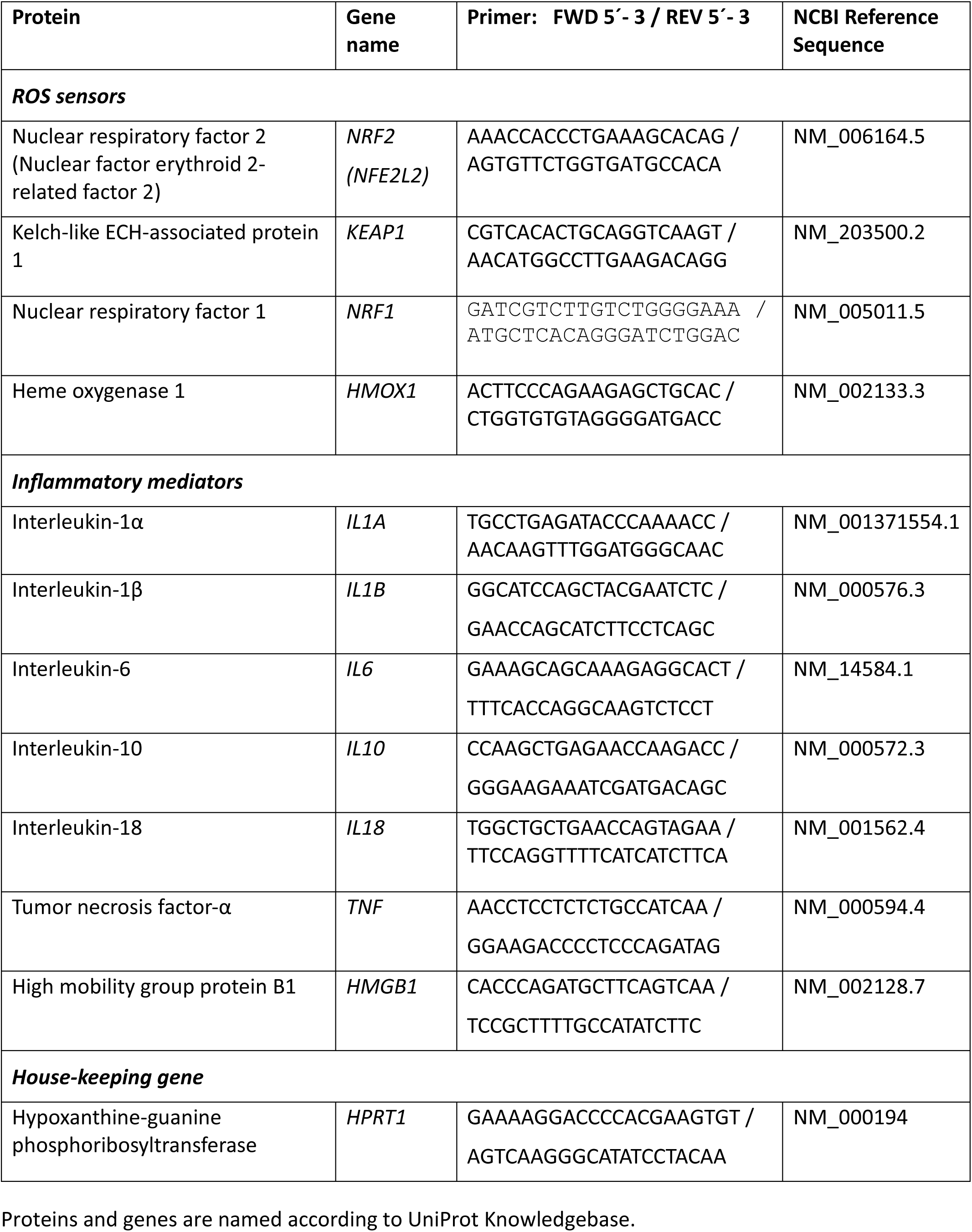
Primers used for quantitative real-time PCR analyses.

### Single muscle fibre collection, typing, and pooling

Approximately 10 mg of muscle biopsy tissue was freeze-dried for 24 h. Biopsies were left in a desiccator in drying pearls (Sigma) at room temperature for 60 min and then stored at-80°C.

Two different methods were used to obtain fibre type-specific pools of muscle fibres.

For the first method, segments of freeze-dried single fibres were collected from each muscle biopsy and placed in 12 ml of 3× SDS denaturing solution (0.125 M Tris-HCI, 10% glycerol, 4% SDS, 4 M urea, 10% 2-mercaptoethanol, and 0.001% Bromophenol Blue, pH 6.8. Collected single fibres were kept at room temperature for 60 min and then stored at-80 °C until dot blotting.

Each single fibre segment was fibre-typed using the dot-blotting method as previously described (Wyckelsma *et al*., 2021). Briefly, a polyvinyldifluorid (PVDF) membrane was activated in 96% ethanol for 120 s and then activated in transfer buffer containing 20% methanol for 120 s. Following activation, 1 ml from each single fibre tube was spotted on to a membrane. Following spotting, membranes were allowed to dry and once dried, they were reactivated in ethanol (120 s) and transfer buffer (120 s). Following reactivation, membranes were washed in Tris-Buffered Saline-Tween (TSBT) and then blocked for 10 min in 5% blocking buffer (Bio-Rad) in TBST. Following blocking, membranes were incubated in primary antibody overnight at 4 °C followed by 2 h at room temperature. One membrane was incubated with myosin heavy chain (MyHC) II (mouse, monoclonal IgG, A4.74, Developmental Studies Hybridoma Bank [DSHB]) and the other membrane incubated with MyHC I (mouse, monoclonal IgM, A4.840, DSHB) diluted 1:200 in blocking buffer in phosphate buffered saline (PBS) (LiCOR Biosciences) 1:1 v/v in 1x TBST.

After washing and incubation with secondary antibody (MyHC II: donkey pAb to mouse IgG (HRP), AbCam, cat n° 98771; MyHC I: goat anti-mouse IgM (heavy chain) Ab (HRP), Thermo Fisher Scientific, cat n° 62-6820), membranes were washed with TBST and coated with chemiluminescent substrate (Clarity Max ECL substrate, Bio-Rad); two membranes were prepared simultaneously from each fibre segment. A Chemidoc MP imaging system (Bio-Rad) was used for membrane imaging.

For the second method, muscle fibres were typed using the THRIFTY protocol (Horwath *et al*., 2022). In brief, the freeze-dried fibres were brought to room temperature on silica gel before handling. Under a stereo microscope, an end of each fibre was cut and mounted onto a microscope slide with a custom grid print, fitting 200 fibres per slide. Once an end of each fibre was mounted to the slides, slides were incubated for 3 min in acetone, followed by 45 min of primary antibody incubation targeting MyHC I (BA-F8-s 1:100, DSHB) and MyHC II (SC-71-s, 1:50, DSHB) diluted in PBS containing 5% normal goat serum (NGS; Thermo Fisher Scientific) and 1% Triton x-100, and followed by 3 x 5 min of PBS wash. Next, 30 min of secondary antibody incubation (Alexa-Fluor goat anti-mouse IgG2b 488, 1:1000 and goat anti-mouse IgG1 647, 1:1000, Thermo Fisher Scientific) in PBS containing 1% NGS and 1% Triton X-100, and another set of 3 x 5 min PBS washes were carried out in dark. Cover glasses were then mounted on the slides using ProLong Gold anti-fade reagent (Thermo Fisher Scientific). Fiber type identification was carried out using a fluorescent microscope (CELENA S Digital Imaging System, Logos Biosystems, Anyang, South Korea).

Single fibre segments from each muscle biopsy identified either as MyHC II or MyHC I with the above-described methods were subsequently pooled into a single tube (∼40 single fibre segments per tube) and stored at-80 °C until used for mass-spectrometry-based proteomics. Pooled single fibre segments were obtained from four participants in the placebo group and five participants in the vitamin group.

### Mass spectrometry-based proteomics

Pooled single fibre segments prepared in 3× SDS denaturing solution were precipitated with cold acetone and centrifuged at 16,000 g for 10 min. Pellets were washed with 80% acetone centrifuged at 16,000 g for 10 min, decanted, air-dried, resuspended in 40 µl of LYSE buffer (PreOmics) in 1.5 ml Eppendorf tubes and heated at 95 °C for 5 min. Samples were then transferred to a Bioruptor sonicator (Diagenode) for 8 min with a 50% duty cycle. Proteolytic digestion was carried out by adding 1 µg of endoproteinase LysC and and 1 µg trypsin per sample. After overnight digestion at 37°C under continuous shaking, the lysate was acidified to a final concentration of 0.1% trifluoro-acetic acid (TFA), and a peptide cleanup was performed using SDB-RPS StageTips as previously described (Kulak *et al*., 2014). The purified peptides were eluted with 60 ul of 80% acetonitrile-1% ammonia and dried in a speed-vac centrifuge for 30 min. Each sample was resuspended in 10 µl of buffer A (0.1% formic acid) and peptide concentration was measured in a Nanodrop 2000 micro-volume spectrophotometer, based on absorbance at 280 nm. Measurements in the mass spectrometer were performed on 300 ug of purified peptides per sample.

Peptides were separated on 50 cm columns of ReproSil-Pur C18-AQ 1.9 μm resin (Dr. Maisch GmbH) packed in-house. The columns were kept at 60 °C using a column oven controlled by the SprayQC software (Scheltema and Mann, 2012). Liquid chromatography was performed on an EASY-nLC 1200 ultra-high-pressure system coupled through a nanoelectrospray source to an Orbitrap Exploris 480 mass spectrometer (all from Thermo Fisher Scientific). Peptides were separated on a nonlinear 45 min gradient of 2–95% buffer A (0.1% formic acid)-buffer B (0.1% formic acid, 80% acetonitrile) at a flow rate of 250 nl/min. Data-independent acquisition was carried out with a master scan of range 350-1400 m/z, followed by 50 DIA scans of 13.7 m/z, with a cycle time of 3 s.

The DIA-NN 1.8.1 (Data-Independent Acquisition by Neural Networks) software (version Apr 14, 2022) was used for the analysis of raw files and peak lists were searched against the human Uniprot FASTA reference proteomes version of 2019 (UP000005640_9606 and corresponding additional file). Library-free search was enabled, and deep learning was used to generate an in silico spectral library from the peptides provided. The output was filtered at 0.01 false discovery rate (FDR). Minimum peptide length was set to 7 and maximum peptide length to 30 amino acids. Cysteine carbamidomethylation was enabled as a fixed modification. The mass spectrometry proteomics data have been deposited to the ProteomeXchange Consortium via the PRIDE partner repository with the dataset identifier PXD055360 (Perez-Riverol *et al*., 2022)

Protein expression values were log2 transformed and proteins with more than 10% missing values were removed using the Perseus software (2.0.11.0), part of the MaxQuant environment (Tyanova *et al*., 2016). The intensity distribution of proteins quantified in the dataset showed the expected large range with the intensity of the abundant and large main myofibrillar proteins (myosins and actin) and myoglobin being seven orders of magnitude larger than the lowest detected proteins (Supplemental Figure 1). The quality of each pooled muscle sample was checked by constructing distribution histograms after subtracting mean values. These histograms showed the expected clear peak and close to normal distribution for each sample except for type 2 fibre samples from a placebo-treated participant, hence type 2 fibre data from this participant were excluded from further analyses (Supplemental Figure 2). Further quality controls with Pearson correlations between measured protein intensities of individual fibre pools in the dataset showed good technical reproducibility between duplicate measurements of each muscle fibre pool (Supplemental Figure 3), and the number of quantified proteins in individual mass spectrometry runs was relatively stable (Supplemental Figure 4).

Thus, the mass spectrometry results presented in the Results are based on pools of type 1 and 2 fibres from five vitamin-treated participants, and pools of type 1 fibres from four and type 2 fibres from three placebo-treated participants. Principal component analyses (a few missing values were replaced from normal distribution), Z-scores, and Volcano plots were generated with the Perseus software; data were exported to Microsoft Excel, where they were organised without further filtering, and figures were subsequently produced with the SigmaPlot software (v11; Systat, Chicago, USA). Relative changes in protein expression induced by the three weeks of SIT were obtained by taking the antilog of log2 post-pre differences; no training-induced change was set to 100%.

### Statistical analysis

Statistical analyses were performed with unpaired *t*-test, paired *t*-test, one-way ANOVA or two-way repeated measures (RM) ANOVA using the SigmaPlot software (v11). *Post hoc* analyses were performed with the Holm–Sidak method when the two-way RM ANOVA showed significant diffierence. The ⍺ level for statistical significance was set to *P* < 0.05. Data are reported as mean ± SD and as individual values.

## RESULTS

### Changes in gene expression after *vs*. before the first and last SIT sessions

Acute SIT-induced changes in gene expression were assessed by measuring mRNA levels in muscle biopsies taken before and one hour after the first and last SIT sessions, i.e. in the untrained and the trained state.

We measured the expression of four genes that are known to respond to ROS challenges in skeletal muscle: *NRF2 (NFE2L2*), encodes a transcription factor that regulates cellular antioxidant capacity and mitochondrial biogenesis; *KEAP1*, encodes a protein that facilitates NRF2 degradation; *NRF1*, encodes a transcription factor that controls genes for key metabolic proteins; *HMOX1*, encodes heme oxygenase, an oxidative stress-induced essential enzyme in heme catabolism (McArdle *et al*., 2004; Merry & Ristow, 2016; Martinez-Canton *et al*., 2024).

In the untrained state, the SIT session induced a significant change in the overall expression of the four ROS-sensitive genes both in the vitamin and the placebo groups (*P* = 0.009 and 0.003, respectively; paired *t*-test, six participants in each group). Notably, the effiect was significantly larger in the placebo than in the vitamin group (*P* < 0.001; two-way RM ANOVA). In the placebo group, the mean expression of all four genes was shifted in the direction expected from an antioxidant effiect of vitamin treatment; that is, mean values increased for *NRF2*, *NRF1* and *HMOX1* and decreased for the negative regulator *KEAP1* (Figure 1*A*; Table 3).

**Figure 1.**
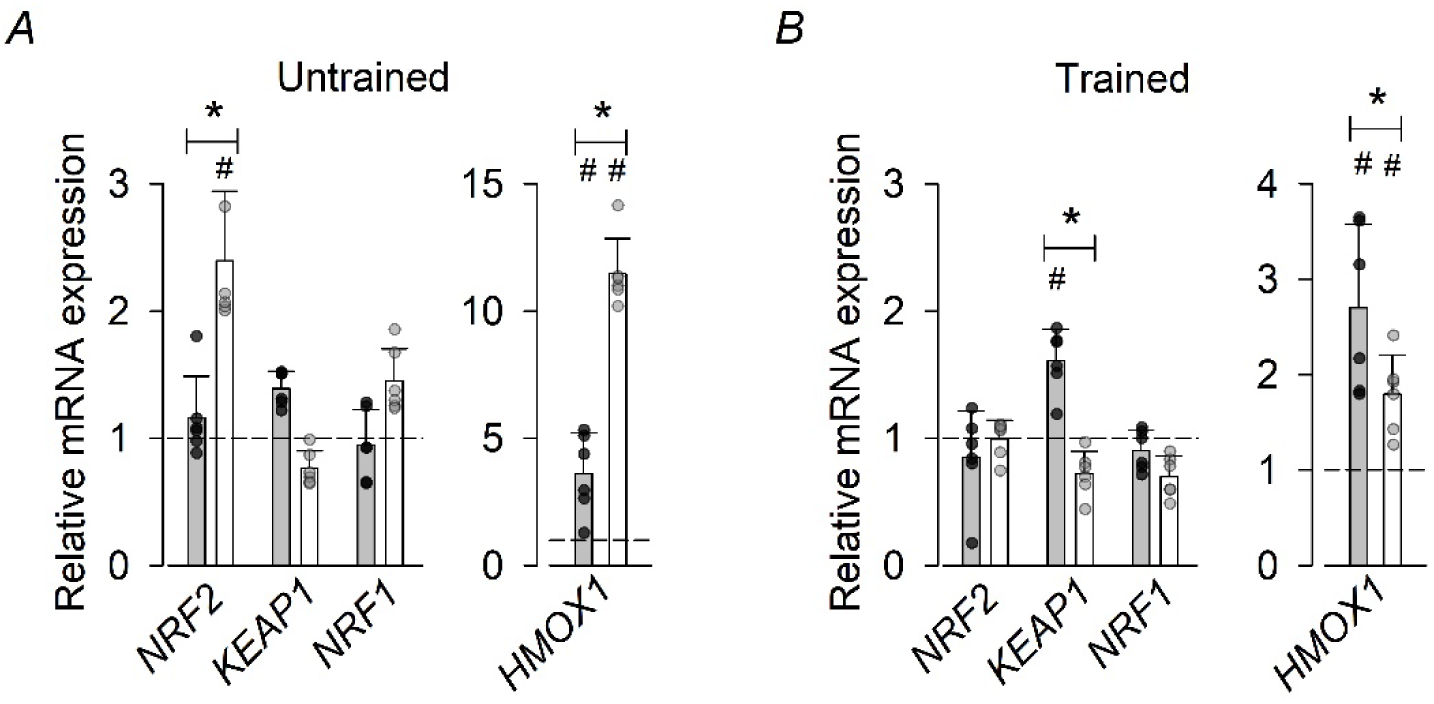
Vitamin treatment blunted the effiect of a SIT session on ROS-sensitive genes in the untrained state. Relative mRNA expression after *vs*. before the first (***A***) and last (***B***) SIT session; no diffierence is set to 1.0. Data are presented as values for individual participants and mean ± SD (n = 6). Vitamin treatment: grey bars and black symbols; placebo treatment: white bars and grey symbols. * *P* < 0.05 between treatments and # *P* < 0.05 after *vs*. before within each group with Holm-Sidak *post hoc* test (precise *P*-values are presented in Table 3). Note the diffierence in y-axis range between ***A*** and ***B*** for *HMOX1*.

**Table 3.** *P*-values for changes in gene expression induced by a SIT session performed either in the untrained or trained states.

| **Gene** | **Untrained** |  |  | **Trained** |  |  |
| --- | --- | --- | --- | --- | --- | --- |
|  | *Vitamin* | *Placebo* | *Vitamin vs. placebo* | *Vitamin* | *Placebo* | *Vitamin vs. placebo* |
| *NRF2* | 0.909 | *0.008* | *0.004* | 0.822 | 0.983 | 0.468 |
| *KEAP1* | 0.880 | 0.568 | 0.130 | *0.012* | 0.496 | *< 0.001* |
| *NRF1* | 0.899 | 0.482 | 0.224 | 0.862 | 0.558 | 0.310 |
| *HMOX1* | *< 0.001* | *< 0.001* | *< 0.001* | *< 0.001* | *0.001* | *< 0.001* |
| *IL1A* | 0.783 | *< 0.001* | *< 0.001* | 0.166 | 0.325 | 0.480 |
| *IL1B* | 0.973 | *< 0.001* | *< 0.001* | *0.006* | *< 0.001* | 0.143 |
| *IL6* | 0.980 | *< 0.001* | *0.004* | 0.432 | *< 0.001* | *< 0.001* |
| *IL10* | 0.989 | *< 0.001* | *< 0.001* | 0.972 | 0.181 | 0.232 |
| *IL18* | 1.000 | 0.776 | 0.772 | 0.437 | 0.988 | *0.004* |
| *TNF* | 0.912 | *< 0.001* | *< 0.001* | 0.987 | 0.083 | *0.047* |
| *HMGB1* | 1.000 | 0.690 | 0.104 | 0.988 | 0.965 | 0.773 |

*P*-values obtained with the Holm-Sidak *post hoc* test; n = 6 participants in each group. *P*-values < 0.05 indicated by italics.

A diffierent pattern emerged in the trained state where a SIT-induced effiect on the ROS-sensitive genes was observed in the vitamin group but not in the placebo group (*P* = 0.009 and 0.616, respectively; paired *t*-test, six participants in each group). Thus, there was still a significant diffierence in the response between the two groups (*P*=0.005; two-way RM ANOVA), but here the response was larger in the vitamin than in the placebo group (Figure 1*B*). *Post hoc* testing showed a significantly larger SIT-induced response in the vitamin group than in the placebo group for *KEAP1* and *HMOX* (Table 3), the latter thus in the opposite direction to the unfatigued state.

ROS challenges are known to trigger inflammatory signalling (Munoz-Canoves *et al*., 2013; Raschke & Eckel, 2013; Peake *et al*., 2015). We measured the acute SIT-induced changes in expression of genes encoding for seven inflammatory mediators: five interleukins (*IL*:s), tumor necrosis factor-α (*TNF*), and high mobility group protein B1 (*HMGB1*).

In the untrained state, both groups showed a highly significant overall up-regulation of the seven inflammatory mediator genes after the SIT session (*P* < 0.001 in both groups; paired *t*-tests, six participants in each group), although vitamin treatment blunted the effiect (*P*<0.001; two-way RM ANOVA) (Figure 2*A*). *Post hoc* testing showed significantly larger SIT-induced effiects on five of the seven genes in the placebo group than in the vitamin group (*IL1A*, *IL1B*, *IL6*, *IL10*, and *TNF*) (Table 3).

**Figure 2.**
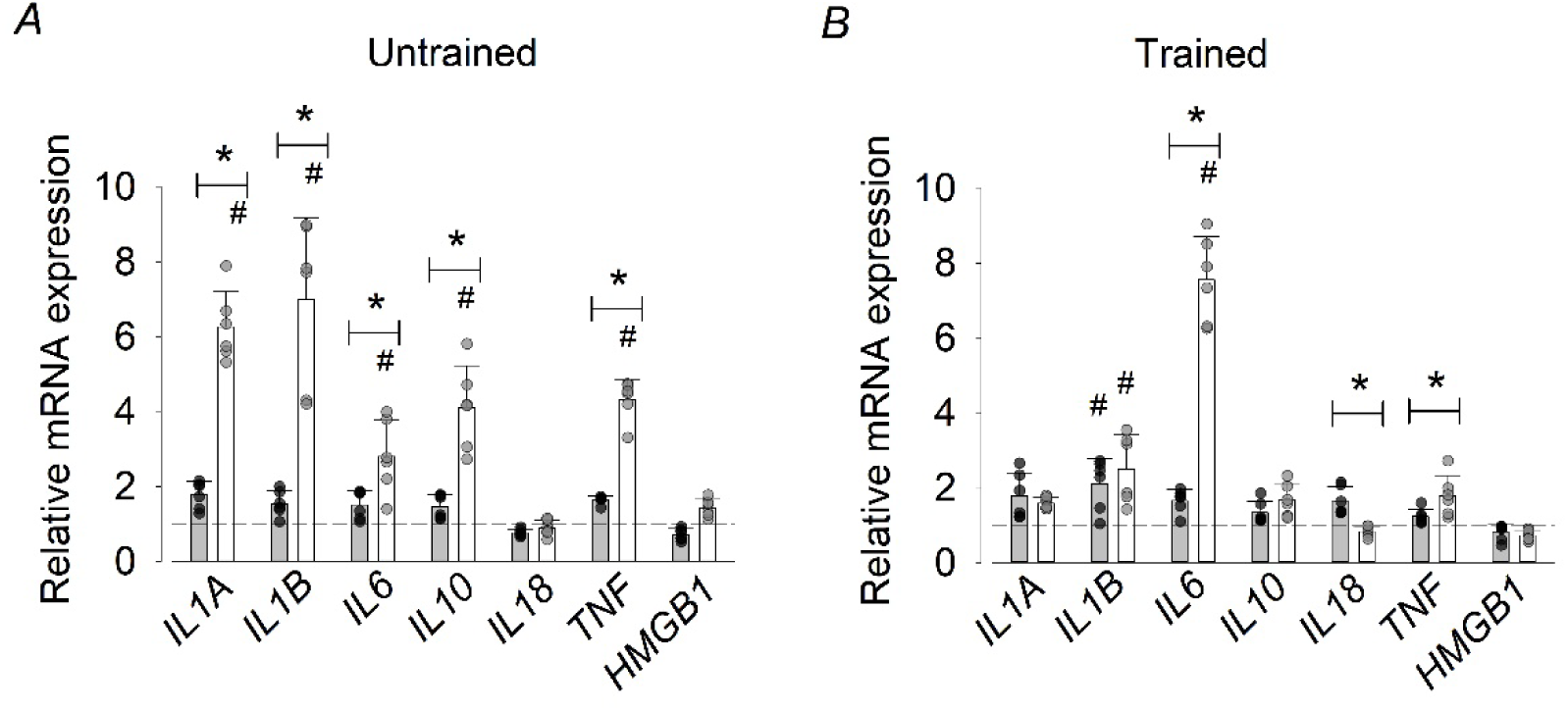
Vitamin treatment blunted the SIT-induced upregulation of genes for inflammatory mediators. Relative mRNA expression after *vs*. before the first (***A***) and last (***B***) SIT session; no diffierence is set to 1.0. Data are presented as values for individual participants and mean ± SD (n = 6). Vitamin treatment: grey bars and black symbols; placebo treatment: white bars and grey symbols. * *P* < 0.05 between treatments and # *P* < 0.05 after *vs*. before within each group with Holm-Sidak *post hoc* test (precise *P*-values are presented in Table 3).

In the trained state, there was still a highly significant SIT-induced overall increase in the expression of the inflammatory mediator genes in both groups (*P* < 0.001 in both groups; paired *t*-tests, six participants in each group). The overall effiect was larger in the placebo than in the vitamin group (*P*<0.001; two-way RM ANOVA) (Figure 2*B*), although the diffierence in SIT-induced effiects was notably smaller in the trained than in the untrained state. At the individual gene level, the only major diffierence between the two groups was a markedly larger response of *IL6* in the placebo than in the vitamin group, although *post hoc* testing also showed significantly diffierent responses between the two groups for *IL18* and *TNF* (Table 3).

### Fibre-type specific proteomics

A total of 2,628 proteins were detected with our proteomics procedures. Proteins were removed from further analyses if they contained more than 10% missing values, which reduced the number of proteins to 1,478 in type 1 fibres and 1,865 in type 2 fibres.

Pooled type 1 fibres (n = 9) contained 90.4 ± 11.2% type 1 myosin (MYH7), whereas the pooled type 2 fibres (n = 8) contained 98.0 ± 2.8% type 2 myosins (MYH1, MYH2, MYH4). Principal component analysis of muscle samples obtained before the three weeks of SIT revealed a clear-cut separation between the two fibre types on the component 1 axis (Figure 3*A*). A Volcano plot of the statistical *P*-value (-log10(*P*)) *vs*. the fold change distribution (log2 diffi (Type 2–Type 1)) showed a clear separation between the two fibre types (Figure 3*B*). To get a crude estimate of the number of protein showing a major diffierence between the two fibre types, we counted the number of proteins with *P* < 0.0001 (-log10(*P*) > 4) and a four-fold diffierence in protein expression (log2 diffi (Type 2–Type 1) >2 or <-2): 21 and 12 proteins displayed a higher expression in type 1 and type 2 fibres, respectively (Figure 3*B*; Table 4). To further illustrate the marked diffierence between the two pools fibres, a Z-scores heatmap of fibre type specific isoforms of SR Ca^2+^ ATPase (SERCA:s/ATP2A:s), calsequestrin (CASQ:s), myosin light chain (MYL:s), and troponin (TNN:s) shows clearly separated expression levels between the two fibre types (Figure 3*C*).

**Figure 3.**
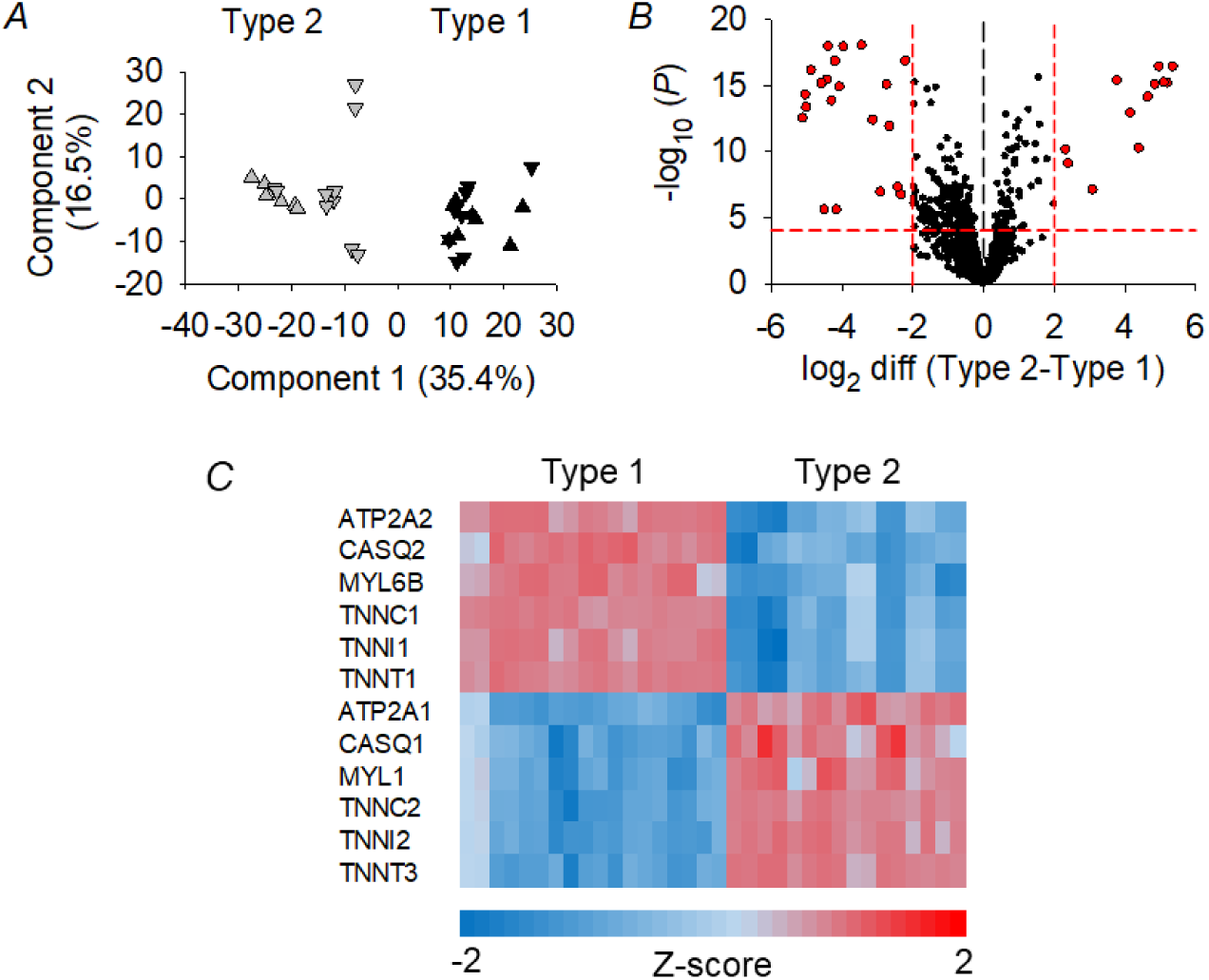
**Proteomics data show clear separation between pooled type 1 and type 2 fibres *A***. Principal component analysis showing the distance between pooled type 1 (▾, vitamin; ▴, placebo) and type 2 (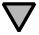, vitamin;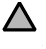, placebo). ***B***. Volcano plot of statistical significance (-log10(*P*)) *vs*. log2 diffierence in protein expression between type 2 and type 1 fibres. Red dashed lines represent levels set to arbitrarily distinguish proteins with expressions being markedly diffierent between the two fibre types (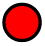). ***C***. Z-scores heatmap illustrating the markedly diffierent expression levels of known fibre type-specific proteins. Data were generated with the Perseus software on duplicates of pooled muscle fibres from five participants in the vitamin group and four (type 1 fibres) or three (type 2 fibres) participants in the placebo group.

**Table 4.** Proteins showing higher expressions before the training period in type 2 and type 1 fibres, respectively.

| | $\log_2$ diff<br>(T2 - T1) | $-\log_{10}(P)$ | Gene | Protein |
| --- | --- | --- | --- | --- |
| **Type 2** | 5.34 | 16.44 | TNNT3 | Isoform of P45378, Troponin T, fast skeletal muscle |
|  | 5.18 | 15.22 | MYH4 | Myosin-4 |
|  | 5.09 | 15.25 | TNNT3 | Troponin T, fast skeletal muscle |
|  | 4.95 | 16.44 | MYH2 | Myosin-2 |
|  | 4.84 | 15.09 | TNNI2 | Troponin I, fast skeletal muscle |
|  | 4.64 | 14.12 | TNNC2 | Troponin C, skeletal muscle |
|  | 4.39 | 10.26 | MYH1 | Myosin-1 |
|  | 4.14 | 12.91 | MYBPC2 | Myosin-binding protein C, fast-type |
|  | 3.77 | 15.40 | MYLPF | Myosin regulatory light chain 2, skeletal muscle |
|  | 3.08 | 7.10 | ACTN3 | Alpha-actinin-3 |
|  | 2.39 | 9.10 | PDLIM7 | Myosin-binding protein C, slow-type |
|  | 2.32 | 10.15 | MYBPC1 | Isoform 2 of PDZ and LIM domain protein 7 |
| **Type 1** | -5.12 | 12.53 | TNNI1 | Troponin I, slow skeletal muscle |
|  | -5.03 | 14.30 | TNNC1 | Troponin C, slow skeletal and cardiac muscles |
|  | -5.00 | 13.35 | MYL3 | Myosin light chain 3 |
|  | -4.87 | 16.16 | TNNT1 | Isoform 2 of Troponin T, slow skeletal muscle |
|  | -4.57 | 15.17 | MYL5 | Myosin light chain 5 |
|  | -4.50 | 5.58 | SHROOM3 | Protein Shroom3 |
|  | -4.43 | 15.43 | MYL2 | Myosin regulatory light chain 2, ventricular/cardiac |
|  | -4.39 | 17.96 | MYH6 | Myosin-6 |
|  | -4.29 | 13.86 | MYL6B | Myosin light chain 6B |
|  | -4.19 | 16.85 | TPM3 | Tropomyosin alpha-3 chain |
|  | -4.16 | 5.59 | TP53BP1 | TP53-binding protein 1 |
|  | -4.07 | 14.90 | ATP2A2 | Isoform 2 of SR/ER calcium ATPase 2 |
|  | -3.95 | 17.95 | MYH7 | Myosin-7 |
|  | -3.45 | 18.06 | MYOZ2 | Myozenin-2 |
|  | -3.13 | 12.40 | USP17L1 | Ubiquitin carboxyl-terminal hydrolase 17-like prot 1 |
|  | -2.91 | 6.94 | DCD | Dermcidin |
|  | -2.74 | 15.08 | DPT | Dermatopontin |
|  | -2.66 | 11.91 | LDHB | L-lactate dehydrogenase B chain |
|  | -2.43 | 7.29 | COX7A2 | Cytochrome c oxidase subunit 7A2, mitochondrial |
|  | -2.33 | 6.73 | COX7A1 | Cytochrome c oxidase subunit 7A1, mitochondrial |
|  | -2.21 | 16.88 | PDLIM1 | PDZ and LIM domain protein 1 |

Listed proteins fulfilled the criteria of $P < 0.0001$ ( $-\log_{10}(P) > 4$ ) and a four-fold difference in protein expression ( $\log_2$ diff (Type 2–Type 1) $> 2$ or $< -2$ ).

#### General response to three weeks of SIT

Volcano plots showing the statistical *P*-value (-log10(*P*)) *vs*. the fold change distribution (log2 diffi (Post-Pre)) illustrate the response to the three weeks of SIT in type 1 fibres from vitamin-and placebo-treated individuals, respectively (Figure 4*A*). The overall picture implies an attenuated training effiect with vitamin treatment. To get a crude estimate of this apparent diffierence, we counted the number of proteins with *P* < 0.01 (-log10(*P*) > 2) and a training effiect larger than ∼40% (log2 diffi (Post-Pre) >0.5 or <-0.7): twice as many proteins fulfilled these criteria in the placebo group (24 proteins) compared to in the vitamin group (12 proteins). The same procedures to assess type 2 fibre data revealed a generally larger training effiect than in type 1 fibres with 107 and 87 proteins exceeding the above criteria in the placebo and the vitamin groups, respectively (Figure 4*B*).

**Figure 4.**
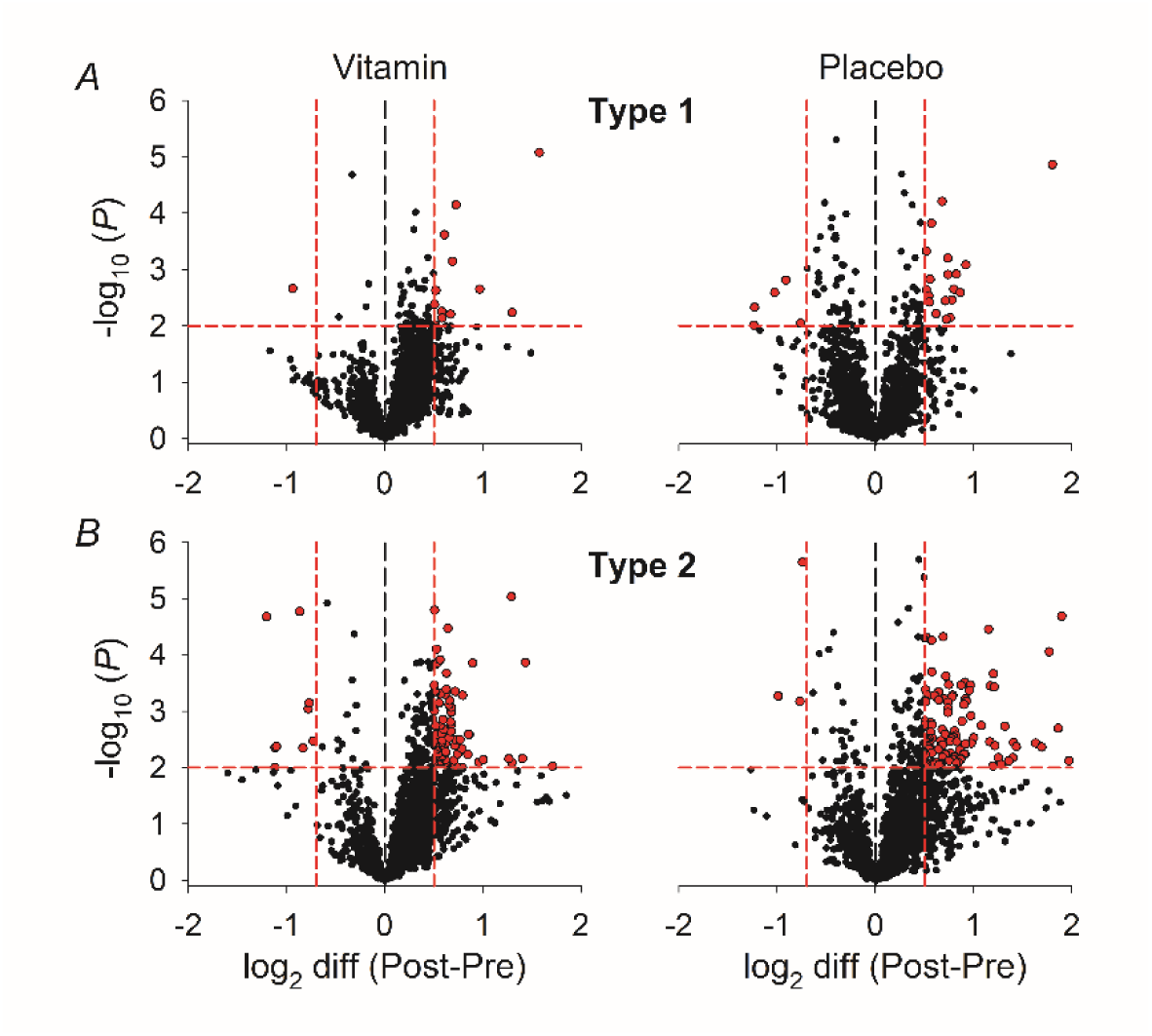
**Three weeks of SIT caused larger proteomic adaptations in type 2 than in type 1 fibres**Volcano plot of statistical significance (-log10(*P*)) *vs*. log2 diffierence in protein expression after vs. before three weeks of SIT in (***A***) type 1 and (***B***) type 2 fibres. Left and right panels show data from vitamin and placebo treated individuals, respectively. Red dashed lines represent levels set to arbitrarily distinguish proteins with expressions being markedly diffierent after *vs*. before the training period (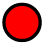). Data were generated with the Perseus software on pooled muscle fibres from five (vitamin, type 1 and 2), four (placebo, type 1) or three (placebo, type 2) participants.

We performed a one-way ANOVA test to assess whether the training response diffiered between the four groups of muscle fibres. For this purpose, the log2 diffierence (Post-Pre) data points in the Volcano plots were converted to relative changes in protein expression, and the statistical testing was performed on their absolute values, i.e. training-induced up-and down-regulations were given the same weight. The results revealed significantly larger training-induced changes in each of the two groups of type 2 fibres (vitamin, 22.8 ± 24.6%; placebo, 25.9 ± 45.6%) than in each of the two type 1 fibre groups (vitamin, 16.8 ± 17.98%; placebo, 17.1 ± 18.4%; *P* < 0.001 for all four combination of groups). Moreover, the training response was significantly larger with placebo than with vitamin treatment in type 2 fibres (*P* = 0.004), whereas there was no significant diffierence between the two groups of type 1 fibres (*P* = 0.789).

In the following, we focus on training-induced changes in three protein groups: ROS-related enzymes, SR Ca^2+^-handling proteins, and mitochondrial proteins involved in aerobic ATP production.

#### ROS-related enzymes

We analysed the effiect of the three weeks of SIT on the expression of eight enzymes critically involved in ROS control in skeletal muscle (Powers & Jackson, 2008): catalse (CAT); glutathione peroxidase 1 (GPX1); peroxiredoxin-1,-3,-5 (PRDX1, PRDX3, PRDX5); cytosolic superoxide dismutase [Cu-Zn] (SOD1); mitochondrial superoxide dismutase [Mn] (SOD2); thioredoxin reductase 1 (TXNRD1).

Figure 5*A* shows the mean training response in type 1 fibres for each of these eight proteins in vitamin *vs*. placebo-treated individuals. Two-way RM ANOVA showed an overall significant diffierence between the two groups (*P* = 0.020); *post hoc* testing revealed a 8.6 ± 8.2% positive training effiect in the vitamin group (*P* = 0.022), a non-significant 6.0 ± 10.6% negative training effiect in the placebo group (*P* = 0.091), and a highly significant diffierence in the the training response between the two groups (*P* < 0.001) (Figure 5B). In type 2 fibres, on the other hand, there was no diffierence in the training response of the ROS-related enzymes (*P* = 0.235; two-way RM ANOVA) (Figure 5C and D).

**Figure 5.**
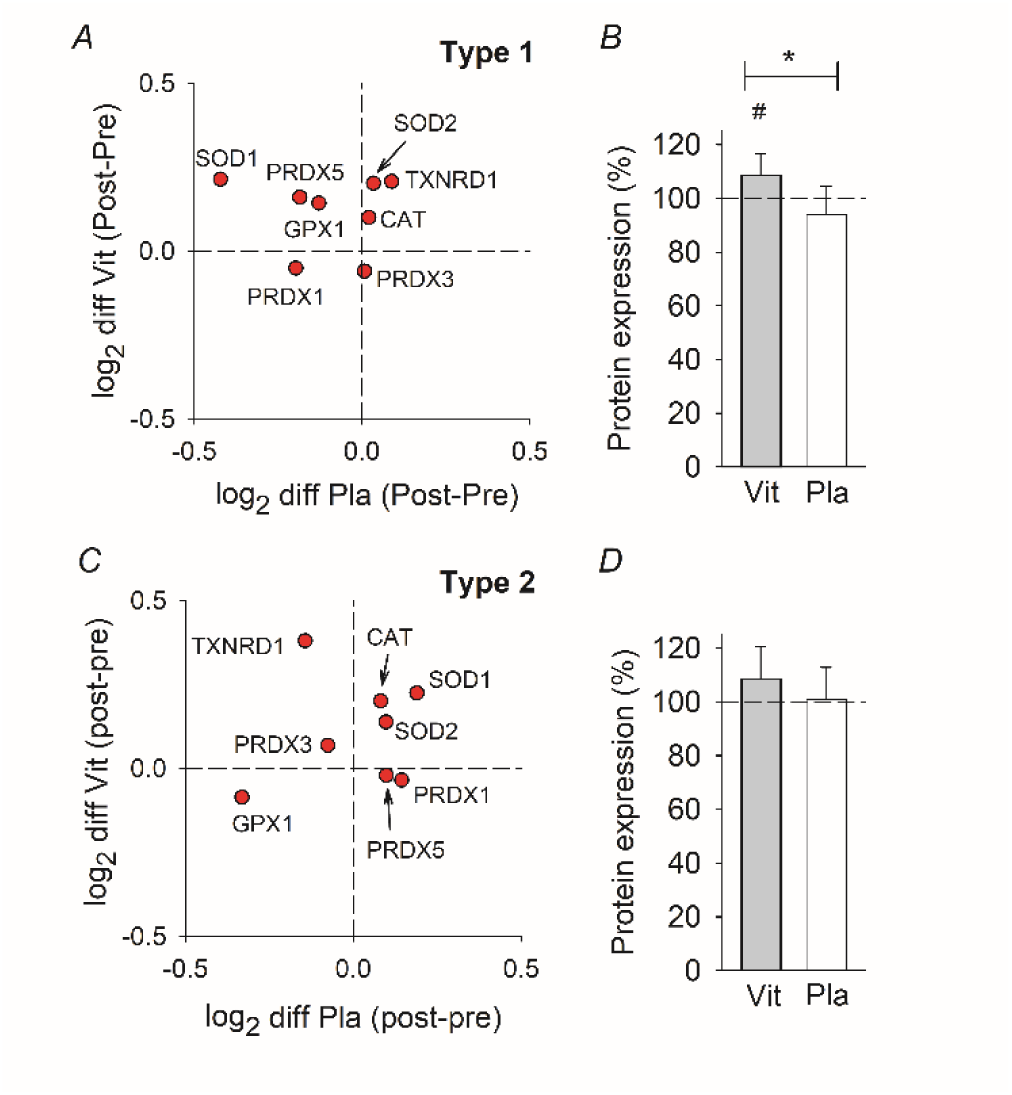
Three weeks of SIT had little impact on the expression of ROS-related enzymes. Training-induced log2 diffierences in expression of eight ROS-related enzymes in vitamin *vs*. placebo treated participants in type 1 (***A***) and type 2 (***C***) fibres. Points (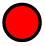) represent the average of five vitamin-treated and four (type 1 fibres) or three (type 2 fibres) placebo-treated individuals. ***B*** and ***D*** show mean (SD) values of the relative overall training-induced changes in expression of the enzymes in type 1 and type 2 fibres, respectively. Grey bars, vitamin treatment; white bars, placebo treatment. Data obtained by antilog of log2 post-pre diffierences; no training effiect was set to 100%. * *P*<0.05 between treatments and # *P*<0.05 within each group with Holm-Sidak *post hoc* test.

#### SR Ca^2+^-handling proteins

Eleven proteins involved in SR Ca^2+^-handling were assessed; four subunits of the dihydropyridine receptor, i.e. the t-tubular volatge sensor (CACNA1S, CACNA2D1, CACNB1, CACNG1); the SR Ca^2+^ release channel (RyR1) and two of its associated proteins, junctophilin 1 (JPH1) and triadin (TRDN); the fast-and slow-twitch isoforms of the SERCA (ATP2A1, ATP2A2); the fast-and slow-twitch isoforms of calsequestrin (CASQ1, CASQ2).

In type 1 fibres, these SR Ca^2+^-handling proteins showed no significant diffierence in the overall training effiect between vitamin and placebo treatment (*P* = 0.830; two-way RM ANOVA) (Figure 6A-B).

**Figure 6.**
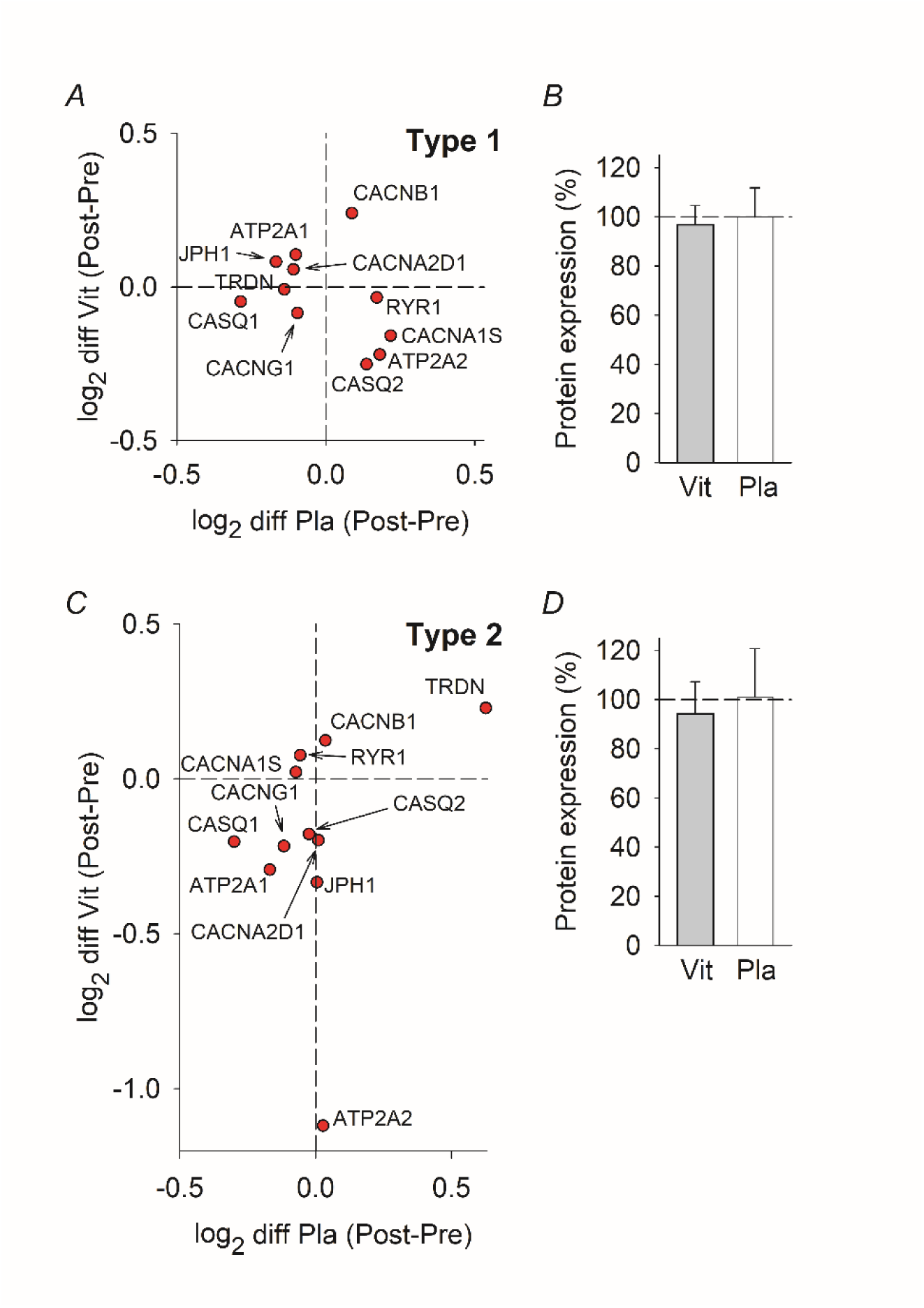
Three weeks of SIT had no significant effiect on the expression of SR Ca^2+^-handling proteins. Training-induced log2 diffierences in expression of eleven SR Ca^2+^-handling proteins in vitamin *vs*. placebo treated participants in type 1 (***A***) and type 2 (***C***) fibres. Points (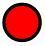) represent the average of five vitamin-treated and four (type 1 fibres) or three (type 2 fibres) placebo-treated individuals. ***B*** and ***D*** show mean (SD) values of the relative overall training-induced changes in protein expression in type 1 and type 2 fibres, respectively. Grey bars, vitamin treatment; white bars, placebo treatment. Data obtained by antilog of log2 post-pre diffierences; no training effiect was set to 100%. SERCA2a (ATP2A2) is not included in *D* because it is expressed at very low levels in type 2 fibres and constitutes a clear outlier in *C*.

In type 2 fibres, SERCA2a (ATP2A2) showed a clear negative training effiect in the vitamin group (Figure 6C). Note that this is the dominant SERCA isoform in type 1 fibres and the expression level is very low in type 2 fibres (Lamboley *et al*., 2014). The remaining ten SR Ca^2+^-handling proteins showed no significant diference in the overall training effiect between the vitamin and placebo groups (*P* = 0.187, two-way RM ANOVA) (Figure 6D).

#### Mitochondrial proteins involved in aerobic ATP production

Proteins related to mitochondrial oxidative capacity were selected with KEGG name filtering using the terms “oxidative phosphorylation” and “citrate cycle”, which resulted in 105 hits in each fibre type. According to MitoCarta 3.0, one of these proteins was not directly involved in aerobic ATP production (PPA2), and six and eight proteins were not localized in mitochondria in type 1 and type 2 fibres, respectively. Thus, analyses were performed on 98 and 96 mitochondrial proteins in type 1 and type 2 fibres, respectively (Supplemental Table 1).

Volcano plots indicated a generally attenuated training effiect with vitamin treatment (Figure 7). To obtain crude measures of this apparent diffierence between the two groups, we counted the number of proteins with *P* < 0.05 (-log10(*P*) > 1.3) and a training effiect larger than ∼20% (log2 diffi (Post-Pre) > 0.25 or <-0.30). In type 1 fibres, one protein fulfilled these criteria in the vitamin group (SDHC down-regulated) compared to eleven proteins (eight up-regulated, three down-regulated) in the placebo group (Figure 7A); corresponding values in type 2 fibres were two up-regulated proteins in the vitamin group *vs*. 18 up-regulated proteins in the placebo group (Fig. 7B).

**Figure 7.**
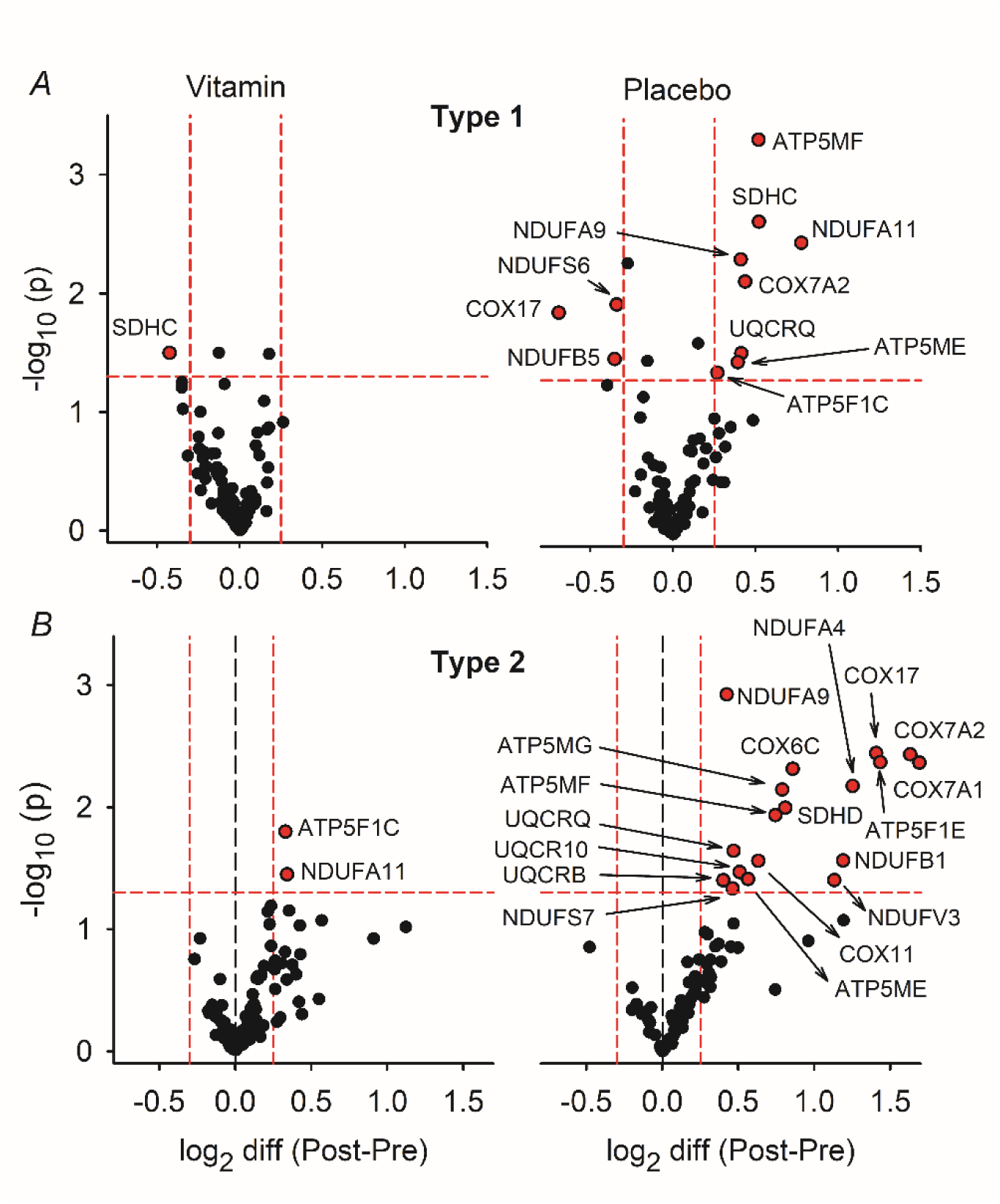
Training-induced effiects on mitochondrial protein expression were larger in the placebo than in the vitamin group. Volcano plot of statistical significance (-log10(*P*)) *vs*. log2 diffierence in mitochondrial protein expression after *vs*. before three weeks of SIT in (***A***) type 1 and (***B***) type 2 fibres. Left and right panels show data from vitamin and placebo treated individuals, respectively. Red dashed lines represent levels set to identify proteins showing large training response (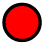 and named). Data were generated with the Perseus software on pooled muscle fibres from five (vitamin, type 1 and 2), four (placebo, type 1) or three (placebo, type 2) participants.

A central aim of the present study was to test whether antioxidant treatment mitigates SIT-induced adaptations towards enhanced muscular aerobic capacity. For this purpose, we used two-way RM ANOVA analysis of the log2 diffi (Post-Pre) data in Figure 7 converted to relative protein expression. For type 1 fibres, the results revealed a significant overall diffierence in training response between the vitamin and the placebo groups (*P* < 0.001). Subsequent *post hoc* testing showed that this was related to a 3.1 ± 8.8% decrease in protein expression in the vitamin group (*P* = 0.018) and a 4.1 ± 15.6% increase in the placebo group (*P* = 0.002). For type 2 fibres, the analysis also revealed an overall diffierence in training response between the two groups (*P* < 0.001); *post hoc* testing showed a 9.8 ± 19.1% increase in the vitamin group (*P* = 0.007) and a 27.9 ± 46.2% increase in the placebo group (*P* < 0.001).

Next, we focused on diffierences in training response between vitamin and placebo treatment on groups of mitochondrial proteins in the pyruvate and tricarboxylic acid cycle (Pyr-TCA) and in complex 1-5 of the electron transport chain (ETC) (Supplemental Table 1). These proteins showed a generally larger training response in type 1 fibres of the placebo than of the vitamin group (*P* < 0.001; two-way RM ANOVA). *Post hoc* testing showed a larger training response in the placebo than in the vitamin group for complex 1 (*P* = 0.018), complex 2 (*P* = 0.033), and complex 3 (*P* = 0.038), whereas no significant diffierences were observed for Pyr-TCA (*P* = 0.996), complex 4 (*P* = 0.117), and complex 5, (*P* = 0.244) (Figure 8A).

**Figure 8.**
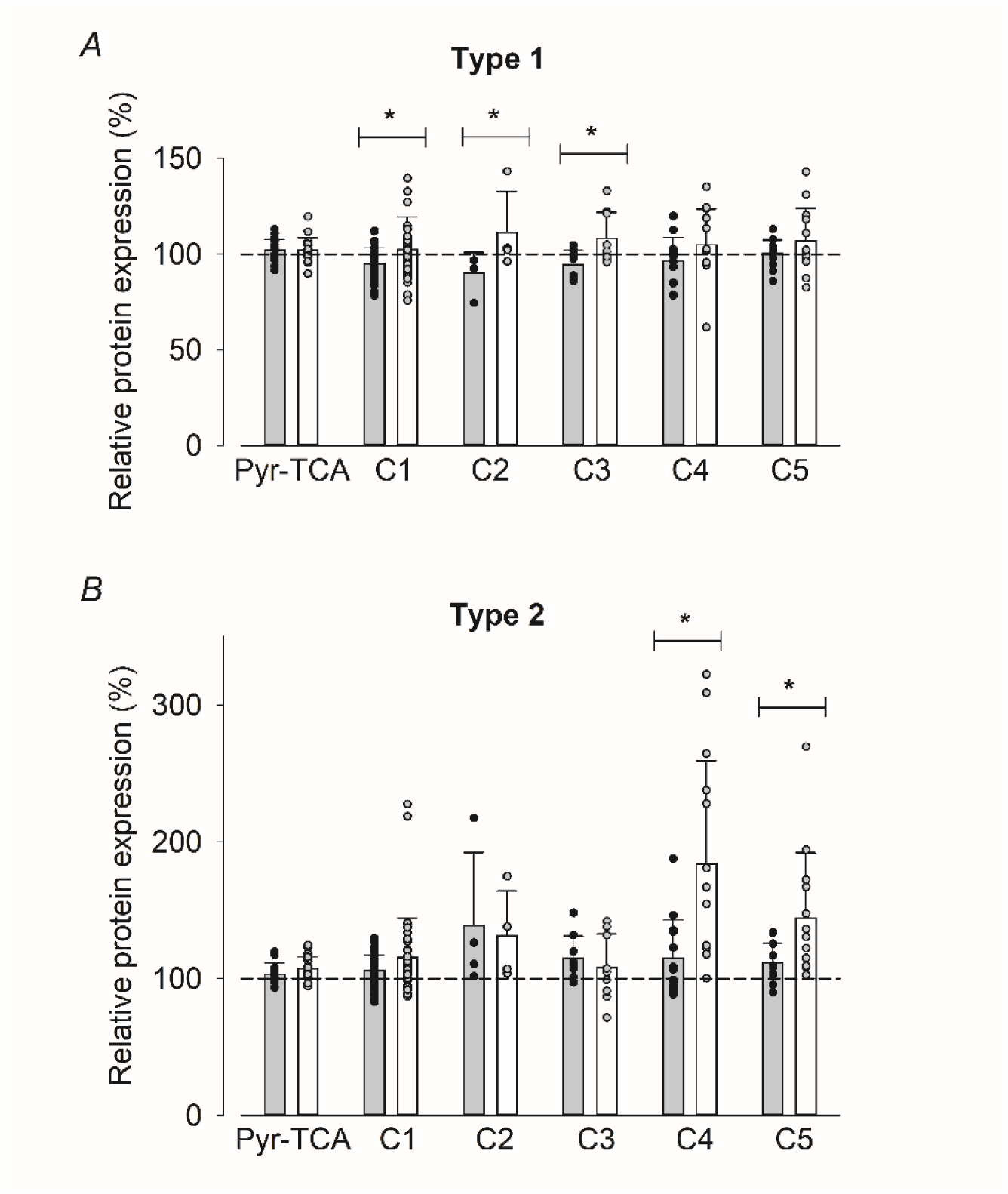
Vitamin treatment blunted the training response in both type 1 and type 2 fibres. Relative expression of proteins in the Pyr-TCA cycle and ETC complexes 1-5 in type 1 (*A*) and type 2 (*B*) fibres. Data obtained by conversion of log2 diffierences after *vs*. before training presented in Figure 7; 100% represents no training effiect. Data presented as mean ± SD and individual vales of each protein. Vitamin treatment: grey bars and black symbols; placebo treatment: white bars and grey symbols. * *P* < 0.05 between treatments within each group with Holm-Sidak *post hoc* test.

In type 2 fibres, the mitochondrial proteins also showed a larger training response in the placebo than in the vitamin group (*P* < 0.001; two-way RM ANOVA). *Post hoc* testing showed larger relative protein expression in the trained state for complex 4 (*P* < 0.001) and complex 5 (*P* < 0.001), whereas no significant diffierences between the two groups were observed for Pyr-TCA (*P* = 0.597), complex 1 (*P* = 0.089), complex 2 (*P* = 0.639), and complex 3, (*P* = 0.548) (Figure 8B).

### Physiology

#### Exercise performance

A standard incremental test to exhaustion on a cycle ergometer was performed in the untrained state (UT; 7 days before the first SIT session) and in the trained state (T; 48 hours after the last SIT session). The three weeks of training resulted in an overall increase in power at exhaustion (*P* < 0.001; two-way RM ANOVA), and *post hoc* tests showed similar increases of ∼6% in the vitamin group (*P* = 0.001) and ∼8% in the placebo group (*P* < 0.001).

Accordingly, there was no diffierence between the two groups either in the untrained (*P* = 0.980) or the trained (*P* = 0.770) state (Figure 9A). Likewise, VO2max was significantly increased after the training period (*P* < 0.001; two-way RM ANOVA); *post hoc* testing revealed similar increases in the vitamin group (∼8%; *P* = 0.001) and in the placebo group (∼6%, *P* = 0.013). Thus, no diffierence between the two groups was detected either in the untrained (*P* = 0.839) or the trained (*P* = 0.909) state (Figure 9B). The heart rate at exhaustion was similar in the untrained and trained state both in the vitamin group (192 ± 6 *vs*. 192 ± 7 min^-1^; *P* = 0.928, paired t-test) and in the placebo group (188 ± 8 *vs*. 188 ± 6 min^-1^; *P* = 0.976).

**Figure 9.**
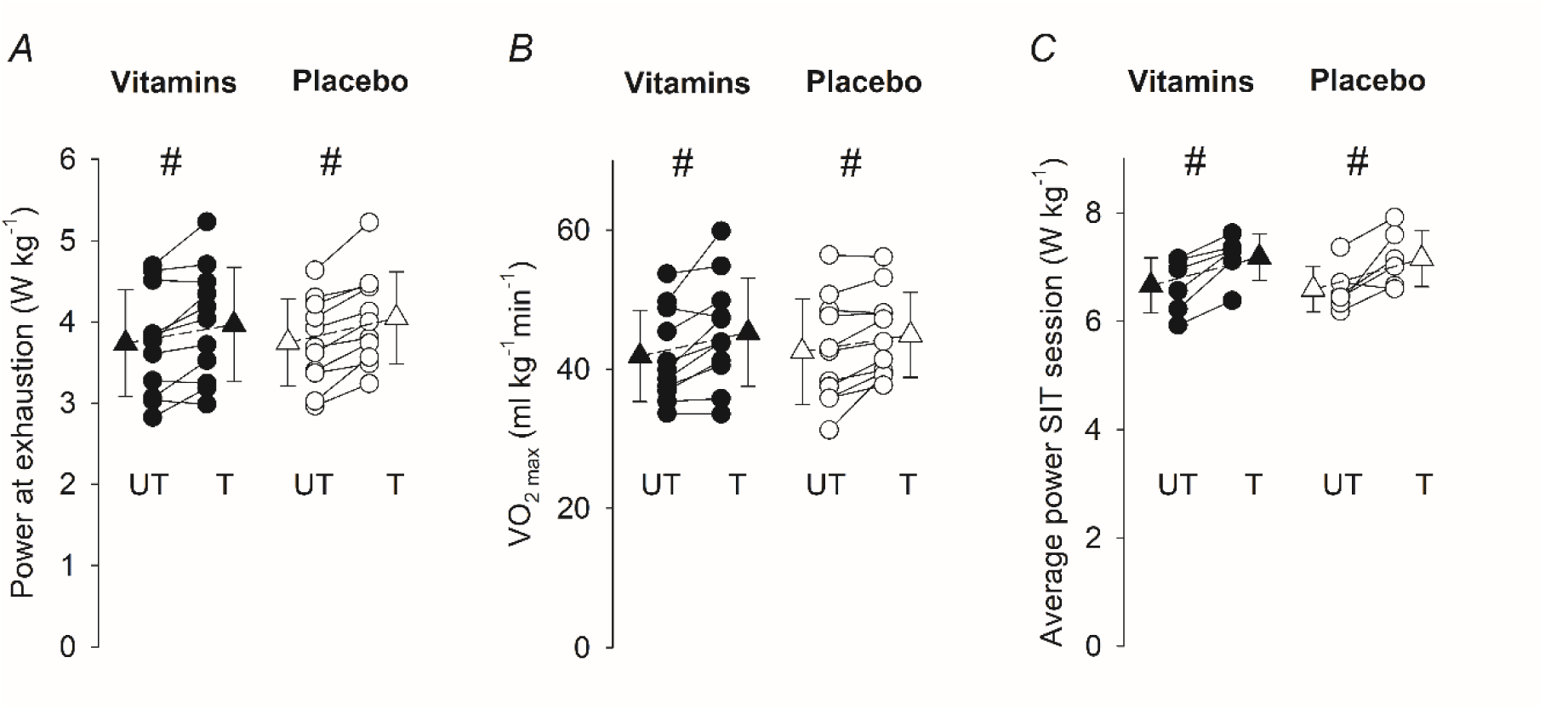
Vitamin treatment did not blunt the training-induced improvement in exercise performance. Power at exhaustion (***A***) and VO2max (***B***) in standard incremental cycling tests performed in the untrained (UT) and trained (T) states. (***C***) Average power during the six Wingate cycling bouts of the first (UT) and last (T) SIT sessions. Data presented as mean (triangles) ± SD and from each participant; n = 11 in *A* and *B*, and n = 6 in *C*. Vitamin, black symbols; placebo, white symbols. # *P* < 0.05 T *vs*. UT with Holm-Sidak *post hoc* test.

The average power during the first and last SIT sessions was measured in a subset of six participants and showed a positive overall training effiect (*P* < 0.001; two-way RM ANOVA), and *post hoc* tests showed similar increases in the vitamin (∼8%, *P* = 0.005) and the placebo (∼9%, *P* = 0.003) groups (Figure 9C). Accordingly, the average power showed no significant diffierence between the two groups either in the untrained (*P* = 0.788) or the trained (*P* = 0.924) state. Blood lactate, measured 5 min after the SIT sessions, was similar in the untrained and trained states both with vitamin (20.1 ± 1,6 *vs*. 18.3 ± 3.4 mM; *P* = 0.228, paired t-test) and placebo (18.0 ± 2.0 *vs*. 19.6 ±2.8 mM; *P* = 0.128) treatment.

Recovery of isometric force was assessed by comparing torques produced before the first (untrained) and before the last (trained) SIT sessions with those produced directly (∼2 min), 1 hour and 24 hours after these sessions. Two-way RM ANOVA analyses showed highly significant (*P* < 0.001) decreases in MVC and 20 Hz torques, and 20 Hz / MVC ratios after the SIT sessions both in the untrained and trained state. *Post hoc* testing revealed significant decreases of all three measures directly and 1 hour after SIT sessions, whereas four out twelve measures remained significantly decreased at 24 hours (Figure 10; Table 5). Importantly, there was no significant diffierence in any of the torque measures between the vitamin and the placebo groups (*P* = 0.556, 0.590, and 0.664 with two-way RM ANOVA for MVC, 20 Hz, and 20Hz/MVC, respectively).

**Figure 10.**
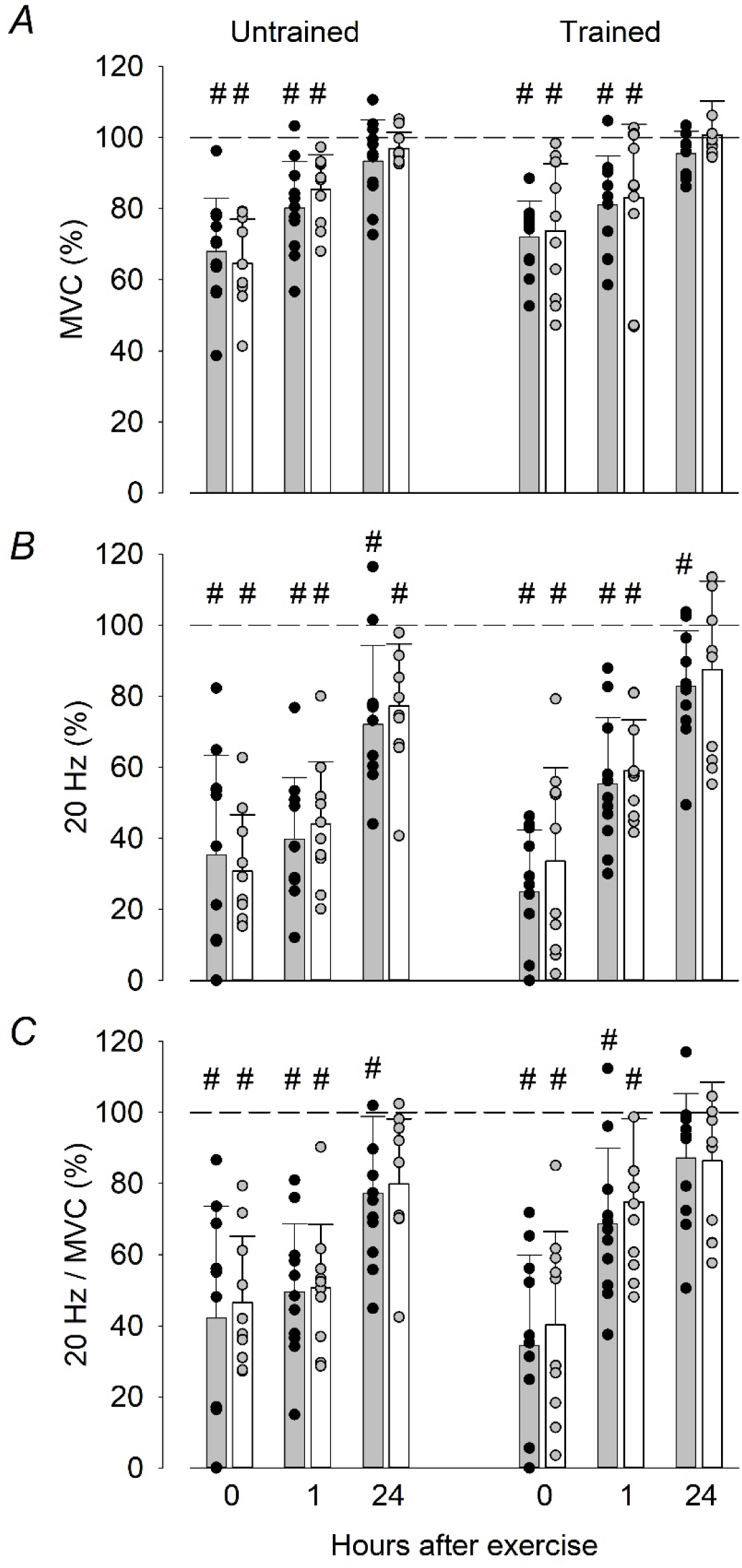
Vitamin treatment did not affiect the extent of PLFFD after SIT sessions. Relative isometric torques at 20 Hz electrical stimulation (***A***) and during MVC (***B***), and the 20 Hz / MVC ratio (***C***) measured during the recovery after the first (Untrained) and last (Trained) SIT sessions. Values are expressed relative to torques measured before the SIT sessions, which were set to 100%. Data presented as mean ± SD and from each participant. Vitamin, grey bars and black symbols (n = 11); placebo, white bars and grey symbols (n = 10). # *P* < 0.05 *vs*. before the SIT session with Holm-Sidak *post hoc* test.

**Table 5.** *P*-values for the reduction in torques after the first (Untrained) and last (Trained) SIT sessions.

|  | Vitamin group (n = 11) |  | Placebo group (n = 10) |  |
| --- | --- | --- | --- | --- |
|  | *Untrained* | *Trained* | *Untrained* | *Trained* |
| **MVC** |  |  |  |  |
| *After* | <0.001 | <0.001 | <0.001 | <0.001 |
| *1 hour* | <0.001 | <0.001 | 0.010 | 0.002 |
| *24 hours* | 0.543 | 0.843 | 0.978 | 0.998 |
| **20 Hz** |  |  |  |  |
| *After* | <0.001 | <0.001 | <0.001 | <0.001 |
| *1 hour* | <0.001 | <0.001 | <0.001 | <0.001 |
| *24 hours* | <0.001 | 0.042 | 0.004 | 0.259 |
| **20Hz/MVC** |  |  |  |  |
| *After* | <0.001 | <0.001 | <0.001 | <0.001 |
| *1 hour* | <0.001 | <0.001 | <0.001 | 0.012 |
| *24 hours* | 0.021 | 0.372 | 0.084 | 0.442 |

*P*-values obtained with Holm-Sidak *post hoc* test. *P*-values < 0.05 indicated by italics.

## Discussion

We tested two major hypotheses: (i) SIT triggers generally larger changes in protein expression in type 2 than in type 1 fibres; (ii) antioxidant supplementation mitigates SIT-induced activations of ROS-sensitive genes that promote muscle fibre adaptations towards increased antioxidant capacity, improved SR Ca^2+^-handling, and enhanced oxidative capacity. In accordance with our first hypothesis, the three weeks of training induced larger changes of the general proteome as well as the expression of mitochondrial proteins in type 2 than in type 1 muscle fibres. In relation to our second hypothesis, the antioxidant treatment inhibited the acute activation of genes encoding for ROS sensors and inflammatory mediators in a manner that might limit beneficial training adaptations. Furthermore, the positive training effiect on mitochondrial protein expression was mitigated by antioxidant supplementation in both fibre types. On the other hand, the three weeks of SIT had no consistent effiect on the expression of selected ROS-related enzymes or SR Ca^2+^ handling proteins, and this was the case both with and without antioxidant supplementation.

After the first SIT session, we observed the expected changes in expression of four genes known to respond to oxidant challenges in the placebo group, and the effiect was blunted in the vitamin group (see Figure 1), which implies a significant antioxidant effiect of the present oral treatment with vitamin C and E (Ristow *et al*., 2009; Paulsen *et al*., 2014; Li *et al*., 2022). A diffierent pattern emerged after the last SIT session where the only expected finding from an oxidant challenge perspective was the larger expression of the negative regulator *KEAP1* in the vitamin than in the placebo group. Thus, the three weeks of training resulted in major changes in the balance between acute SIT-induced ROS production and defence, and it appeared likely that this would involve increased abundance of enzymes critically involved in the cellular ROS defence (Jordan *et al*., 2021; Martinez-Canton *et al*., 2024; Powers *et al*., 2024). However, our proteomics data showed relatively minor changes in abundance of eight ROS-controlling enzymes (see Figure 5). Moreover, the only statistically significant training-induced adaptation at the group level was an ∼11% increase in expression in type 1 fibres of the vitamin group, thus opposite to what would be expected from an adaptative response triggered by oxidant challenge. A simplistic interpretation of these findings is that main ROS-related adaptations affiected the production rather than the defence side of the redox balance, but further studies are required to establish whether this is the case.

After the first SIT session, genes encoding for cytokines showed a marked up-regulation in the placebo group, and this response was mitigated in the vitamin group (see Figure 2A). Thus, acute inflammatory-related signalling can be considered as a normal ROS-dependent and possibly beneficial response to physiological stress, such as a SIT session, which stands in sharp contrast to the prolonged oxidant and inflammatory signalling observed with excessive training (“overtraining”) and in inflammatory disorders that is associated with functional impairments and muscle wasting (Munoz-Canoves *et al*., 2013; Raschke & Eckel, 2013; Peake *et al*., 2015; Cheng *et al*., 2020).

The acute SIT-induced effiect on cytokine gene expression was generally smaller in the trained than in the untrained state, and the only remaining clear-cut diffierence between the two groups was an almost eight-fold increase in *IL6* expression in the placebo group (see Figure 2B). Notably, transiently increased IL6 production in muscle fibres is associated with beneficial autocrine and paracrine effiects, whereas prolonged systemic production is linked to deleterious effiects, such as muscle wasting (Munoz-Canoves *et al*., 2013).

Altered SR Ca^2+^ handling causing moderate increases SR Ca^2+^ leak and free cytosolic [Ca^2+^] in resting muscle fibres can trigger mitochondrial biogenesis and hence lead to increased endurance (Ojuka *et al*., 2002; Wright *et al*., 2007; Aydin *et al*., 2008; Ivarsson *et al*., 2019). ROS challenges can affiect the RyR1 protein complex that controls SR Ca^2+^ release by causing dissociation of the channel stabilizing protein FKBP12 (Bellinger *et al*., 2008a).

Moreover, fragmentation of RyR1 after a demanding SIT session was first observed in muscle biopsies of young men, and parallel experiments on isolated mouse muscle indicated that the fragmentation was ROS-dependent (Place *et al*., 2015). Subsequent studies show that this acute RyR1 fragmentation also occurs in muscles of elderly men (Wyckelsma *et al*., 2020), is restricted to type 2 fibres (Tripp *et al*., 2022), and its prevalence is decreased after three weeks of SIT (Schlittler *et al*., 2019; Wyckelsma *et al*., 2020). In addition, five weeks of high-intensity interval training has been shown to decrease the expression of DHPR subunits in vastus lateralis muscles (Hostrup *et al*., 2022). Based on the findings outlined above, we expected clear-cut ROS-dependent adaptions in the expression of SR Ca^2+^-handling proteins at the end of the three weeks of SIT. However, no consistent training induced changes in expression of SR Ca^2+^-handling proteins were observed either in type 1 or type 2 fibres, and this was the case both for the vitamin and the placebo group (see Figure 6). The only exception was a clear negative training effiect in the expression of SERCA2a in type 2 fibres of vitamin-treated participants, which would be of very limited functional importance since it is the dominant SERCA isoform in type 1 fibres and the expression level is very low in type 2 fibres (Lamboley *et al*., 2014). Thus, SIT-induced changes in SR Ca^2+^-handling seem related to post-translational protein modifications, and these modifications are then not accompanied by any consistent adaptive changes in the abundance of proteins with key roles in SR Ca^2+^ release, Ca^2+^ buffiering or active Ca^2+^ uptake.

In accordance with our hypotheses, the three weeks of SIT resulted in increased expression of proteins involved in mitochondrial energy production, and the effiect was larger in type 2 than in type 1 fibres and blunted by vitamin treatment. The greater effiect in type 2 fibres can be explained by these fibres being recruited to a large extent during SIT (Vøllestad & Blom, 1985) and thereby being exposed to a greater adaptation-promoting metabolic stress than type 1 fibres (Henriksson & Reitman, 1976; Dudley *et al*., 1982; MacInnis & Gibala, 2017). In line with this reasoning, larger training-induced proteomic changes in type 1 than in type 2 fibres were observed after 12 weeks of moderate-intensity endurance training, where type 2 fibres are less recruited than during SIT (Deshmukh *et al*., 2021). A recent study on fibre-type specific proteomics reports no increases in abundance of ETC complex 1-5 or TCA cycle proteins after 8 weeks of SIT (Reisman *et al*., 2024a, see their Supplemental Figure 2). In relation to our findings, the lack of a positive training effiect observed by Reisman *et al*. is in reasonable agreement with the minor (∼4%) increase in mitochondrial protein expression in type 1 fibres of the placebo group. In relation to type 2 fibres, on the other hand, the lack of effiect observed by Reisman *et al*. is in sharp contrast to our large (average

∼28%) SIT-induced increase in mitochondrial protein expression, as well as previously presented high-intensity exercise-induced increases in mitochondrial protein concentration in type 2 fibres (Henriksson & Reitman, 1976; Dudley *et al*., 1982; Tan *et al*., 2018; Skelly *et al*., 2021). The participants in the study by Reisman *et al*. (2024) had higher oxidative capacity before the training period than our participants, as judged from markedly higher mean values during incremental cycling tests: VO2max, 50.8 vs. 42.5 ml kg^-1^ min^-1^; maximal power, 4.2 vs. 3.7 W kg^-1^. Our participants produced markedly higher power during SIT sessions than the maximal power reached at the end of the incremental cycling test (see Figure 9), which implies large recruitment of type 2 fibres during SIT sessions; corresponding data were not presented by Reisman et al. (2024). Thus, the lack of a SIT-induced increase in mitochondrial protein abundance in the study of Reisman et al. might, for instance, be explained by limited type 2 fibre recruitment and a metabolic stress not being large enough to trigger adaptations in individuals with an already high oxidative capacity.

Our results show a significant blunting of the training-induced increase in mitochondrial protein expression with vitamin treatment in both fibre types, which adds to many previous studies showing an opposing effiect of antioxidant supplementation on beneficial adaptations to endurance training (Gomez-Cabrera *et al*., 2008; Ristow *et al*., 2009; Strobel *et al*., 2011; Paulsen *et al*., 2014). Intriguingly, the effiect of vitamin treatment on the training response diffiered between fibre types in that complex 1-3 proteins were significantly lowered in type 1 fibres, whereas complex 4-5 proteins were significantly lowered in type 2 fibres (see Figure 8). The mechanisms underlying this diffierence in antioxidant response between the two fibre types and its physiological significance require further studies.

The three weeks of SIT increased VO2max and the power at the end of the incremental cycling test as well as the average power during SIT sessions to the same extent in vitamin-and placebo-treated participants (see Figure 9). VO2max represents the highest rate at which O2 can be taken up and utilized during intense physical exercise (Bassett & Howley, 2000). It depends on the integrated ability of pulmonary, cardiovascular, and skeletal muscle systems to take up, transport, and utilize O2, which mainly occurs in the mitochondria of contracting muscles (Dominelli *et al*., 2021; Gibala & MacInnis, 2022). Thus, any step(s) in this functional cascade can be rate limiting for VO2max. Classically, VO2max is considered to be limited by factors that determine O2 delivery to working muscles (Bassett & Howley, 2000; Lundby *et al*., 2017), although the O2-utilizing capacity of mitochondria in exercising muscles might set the limit in untrained individuals (Giffiord *et al*., 2016). In a recent study on healthy men and women, six weeks of SIT increased VO2max by ∼10% and this could be explained by improved O2 delivery due to increases in both the total circulation blood volume and the left ventricular volume (Eriksson *et al*., 2024). Accordingly, our results fit with O2 delivery to the working muscles being limiting, since vitamin-and placebo-treated participants showed similar training-induced improvements in exercise performance, including increased VO2max, despite blunted increases in mitochondrial protein abundance in muscles of vitamin-treated individuals. Importantly, this implies ROS-independent mechanisms underlying the training-induced improvement of O2 delivery, whereas the increase in mitochondrial protein abundance was triggered by ROS-dependent mechanisms. Nevertheless, the SIT-induced increase in mitochondrial content in muscle will improve endurance exercise performance, for instance, due to decreased lactate production at a given submaximal power output, hence limiting the depletion of muscular glycogen stores (Coyle *et al*., 1988; Bassett & Howley, 2000).

Previous studies show a prolonged depression of isometric knee extension torque after SIT sessions especially at low stimulation frequencies, that is, muscles entered a state of PLFFD (Place *et al*., 2015; Schlittler *et al*., 2019; Wyckelsma *et al*., 2020). The present results show PLFFD after the first and last SIT sessions (see Figure 10). Notably, the recovery of torques and extent of PLFFD were not affiected by vitamin treatment, which agrees with previous results obtained in elderly individuals (Wyckelsma *et al*., 2020). Experiments on isolated muscle fibres have revealed two major mechanisms underlying PLFFD after fatiguing contractions: decreased SR Ca^2+^ release and reduced myofibrillar Ca^2+^ sensitivity (Allen *et al*., 2008; Richards *et al*., 2025). The relative importance of these two mechanisms has been shown to be ROS-dependent (Bruton *et al*., 2008; Cheng *et al*., 2016), and addition of antioxidants has been shown to shift the dominant mechanism from decreased SR Ca^2+^ release to reduced myofibrillar Ca^2+^ sensitivity without affiecting the extent of PLFFD (Cheng *et al*., 2015). Thus, the present results can be explained by vitamin treatment moving the cause of PLFFD towards reduced myofibrillar Ca^2+^ sensitivity without affiecting its magnitude.

In conclusion, in the untrained state, SIT activates ROS-dependent signalling that is translated into proteomic changes towards improved muscular aerobic capacity. This positive training response is greater in type 2 than in type 1 muscle fibres, which reflects large recruitment of type 2 fibres during SIT and can explain why SIT is an effiective complement to endurance training performed at lower intensities. Antioxidant supplementation does not blunt the SIT-induced improvement in maximal exercise performance, which implies ROS-independent triggering of cardiovascular adaptations towards improved O2 delivery to muscle. Nevertheless, a SIT-induced increase in muscular aerobic capacity would improve performance during submaximal exercise.

## Data availability statement

The mass spectrometry proteomics data have been deposited to the ProteomeXchange Consortium via the PRIDE partner repository with the dataset identifier PXD055360 (Perez-Riverol *et al*., 2022). Other data that support the findings of this study are available from the corresponding author upon reasonable request.

## Author contributions

VLW, SK, MB, HW and TV were responsible for the study conception and design. VLW, MM, NE, AS, MP, SG, SE, WA, HW and TV were responsible for data acquisition, as well as analysis and interpretation. VLW, MM, SK, MB, MP, SG, SE, DCA, HW and TV were responsible for drafting and revising the article. All authors approved final version of the manuscript and agree to be accountable for all aspects of the work

## Acknowledgements

The mouse, monoclonal IgG, A4.74 and mouse, monoclonal IgM, A4.840 antibodies were obtained from the Developmental Studies Hybridoma Bank, created by the NICHD of the NIH and maintained at The University of Iowa, Department of Biology, Iowa City, IA 52242.

## References

Aguilo A, Tauler P, Sureda A, Cases N, Tur J & Pons A. (2007). Antioxidant diet supplementation enhances aerobic performance in amateur sportsmen. J Sports Sci 25, 1203–1210.

Allen DG, Lamb GD & Westerblad H. (2008). Skeletal muscle fatigue: cellular mechanisms. Physiol Rev 88, 287–332.

Aydin J, Shabalina IG, Place N, Reiken S, Zhang SJ, Bellinger AM, Nedergaard J, Cannon B, Marks AR, Bruton JD & Westerblad H. (2008). Nonshivering thermogenesis protects against defective calcium handling in muscle. FASEB J 22, 3919–3924.

Bassett DR, Jr. & Howley ET. (2000). Limiting factors for maximum oxygen uptake and determinants of endurance performance. Med Sci Sports Exerc 32, 70–84.

Bellinger AM, Mongillo M & Marks AR. (2008a). Stressed out: the skeletal muscle ryanodine receptor as a target of stress. J Clin Invest 118, 445–453.

Bellinger AM, Reiken S, Dura M, Murphy PW, Deng SX, Landry DW, Nieman D, Lehnart SE, Samaru M, Lacampagne A & Marks AR. (2008b). Remodeling of ryanodine receptor complex causes “leaky” channels: a molecular mechanism for decreased exercise capacity. Proc Natl Acad Sci U S A 105, 2198–2202.

Bruton JD, Place N, Yamada T, Silva JP, Andrade FH, Dahlstedt AJ, Zhang SJ, Katz A, Larsson NG & Westerblad H. (2008). Reactive oxygen species and fatigue-induced prolonged low-frequency force depression in skeletal muscle fibres of rats, mice and SOD2 overexpressing mice. J Physiol 586, 175–184.

Cheng AJ, Bruton JD, Lanner JT & Westerblad H. (2015). Antioxidant treatments do not improve force recovery after fatiguing stimulation of mouse skeletal muscle fibres. J Physiol 593, 457–472.

Cheng AJ, Jude B & Lanner JT. (2020). Intramuscular mechanisms of overtraining. Redox Biol 35, 101480.

Cheng AJ, Yamada T, Rassier D, Andersson DC, Westerblad H & Lanner JT. (2016). Reactive oxygen/nitrogen species and contractile function in skeletal muscle during fatigue and recovery. J Physiol 594, 5149–5160.

Coyle EF, Coggan AR, Hopper MK & Walters TJ. (1988). Determinants of endurance in well-trained cyclists. J Appl Physiol 64, 2622–2630.

Deshmukh AS, Steenberg DE, Hostrup M, Birk JB, Larsen JK, Santos A, Kjobsted R, Hingst JR, Scheele CC, Murgia M, Kiens B, Richter EA, Mann M & Wojtaszewski JFP. (2021). Deep muscle-proteomic analysis of freeze-dried human muscle biopsies reveals fiber type-specific adaptations to exercise training. Nat Commun 12, 304.

Dominelli PB, Wiggins CC, Roy TK, Secomb TW, Curry TB & Joyner MJ. (2021). The oxygen cascade during exercise in health and disease. Mayo Clin Proc 96, 1017–1032.

Dudley GA, Abraham WM & Terjung RL. (1982). Influence of exercise intensity and duration on biochemical adaptations in skeletal muscle. J Appl Physiol Respir Environ Exerc Physiol 53, 844–850.

Eriksson LMJ, Hedman K, Åstrom-Aneq M, Nylander E, Bouma K, Mandic M, Gustafsson T & Rullman E. (2024). Evidence of left ventricular cardiac remodeling after 6 weeks of sprint interval training. Scand J Med Sci Sports 34, e70007.

Flockhart M, Nilsson LC, Tillqvist EN, Vinge F, Millbert F, Lannerström J, Nilsson PH, Samyn D, Apro W, Sundqvist ML & Larsen FJ. (2023). Glucosinolate-rich broccoli sprouts protect against oxidative stress and improve adaptations to intense exercise training. Redox Biol 67, 102873.

Gibala MJ & MacInnis MJ. (2022). Physiological basis of brief, intense interval training to enhance maximal oxygen uptake: a mini-review. Am J Physiol Cell Physiol 323, C1410–C1416.

Giffiord JR, Garten RS, Nelson AD, Trinity JD, Layec G, Witman MA, Weavil JC, Mangum T, Hart C, Etheredge C, Jessop J, Bledsoe A, Morgan DE, Wray DW, Rossman MJ & Richardson RS. (2016). Symmorphosis and skeletal muscle VO_2max_: *in vivo* and *in vitro* measures reveal diffiering constraints in the exercise-trained and untrained human. J Physiol 594, 1741–1751.

Gomez-Cabrera MC, Domenech E, Romagnoli M, Arduini A, Borras C, Pallardo FV, Sastre J & Vina J. (2008). Oral administration of vitamin C decreases muscle mitochondrial biogenesis and hampers training-induced adaptations in endurance performance. Am J Clin Nutr 87, 142–149.

Henriksson J & Reitman JS. (1976). Quantitative measures of enzyme activities in type I and type II muscle fibres of man after training. Acta Physiol Scand 97, 392–397.

Horwath O, Edman S, Andersson A, Larsen FJ & Apro W. (2022). THRIFTY: a novel high-throughput method for rapid fibre type identification of isolated skeletal muscle fibres. J Physiol 600, 4421–4438.

Hostrup M, Lemminger AK, Stocks B, Gonzalez-Franquesa A, Larsen JK, Quesada JP, Thomassen M, Weinert BT, Bangsbo J & Deshmukh AS. (2022). High-intensity interval training remodels the proteome and acetylome of human skeletal muscle. Elife 11, e69802.

Ivarsson N, Mattsson CM, Cheng AJ, Bruton JD, Ekblom B, Lanner JT & Westerblad H. (2019). SR Ca^2+^ leak in skeletal muscle fibers acts as an intracellular signal to increase fatigue resistance. J Gen Physiol 151, 567–577.

Jordan AC, Perry CGR & Cheng AJ. (2021). Promoting a pro-oxidant state in skeletal muscle: Potential dietary, environmental, and exercise interventions for enhancing endurance-training adaptations. Free Radic Biol Med 176, 189–202.

Kulak NA, Pichler G, Paron I, Nagaraj N & Mann M. (2014). Minimal, encapsulated proteomic-sample processing applied to copy-number estimation in eukaryotic cells. Nat Methods 11, 319–324.

Lamboley CR, Murphy RM, McKenna MJ & Lamb GD. (2014). Sarcoplasmic reticulum Ca^2+^ uptake and leak properties, and SERCA isoform expression, in type I and type II fibres of human skeletal muscle. J Physiol 592, 1381–1395.

Li S, Fasipe B & Laher I. (2022). Potential harms of supplementation with high doses of antioxidants in athletes. J Exerc Sci Fit 20, 269–275.

Lundby C, Montero D & Joyner M. (2017). Biology of VO2max: looking under the physiology lamp. Acta Physiol (Oxf) 220, 218–228.

MacInnis MJ & Gibala MJ. (2017). Physiological adaptations to interval training and the role of exercise intensity. J Physiol 595, 2915–2930.

Magistris MR, Kohler A, Pizzolato G, Morris MA, Baroffio A, Bernheim L & Bader CR. (1998). Needle muscle biopsy in the investigation of neuromuscular disorders. Muscle Nerve 21, 194–200.

Margaritelis NV, Theodorou AA, Paschalis V, Veskoukis AS, Dipla K, Zafeiridis A, Panayiotou G, Vrabas IS, Kyparos A & Nikolaidis MG. (2018). Adaptations to endurance training depend on exercise-induced oxidative stress: exploiting redox interindividual variability. Acta Physiol (Oxf) 222, DOI: 10.1111/apha.12898.

Martinez-Canton M, Galvan-Alvarez V, Martin-Rincon M, Calbet JAL & Gallego-Selles A. (2024). Unlocking peak performance: The role of Nrf2 in enhancing exercise outcomes and training adaptation in humans. Free Radic Biol Med 224, 168–181.

McArdle F, Spiers S, Aldemir H, Vasilaki A, Beaver A, Iwanejko L, McArdle A & Jackson MJ. (2004). Preconditioning of skeletal muscle against contraction-induced damage: the role of adaptations to oxidants in mice. J Physiol 561, 233–244.

Merry TL & Ristow M. (2016). Nuclear factor erythroid-derived 2-like 2 (NFE2L2, Nrf2) mediates exercise-induced mitochondrial biogenesis and the anti-oxidant response in mice. J Physiol 594, 5195–5207.

Munoz-Canoves P, Scheele C, Pedersen BK & Serrano AL. (2013). Interleukin-6 myokine signaling in skeletal muscle: a double-edged sword? Febs J 280, 4131–4148.

Ojuka EO, Jones TE, Han DH, Chen M, Wamhoffi BR, Sturek M & Holloszy JO. (2002). Intermittent increases in cytosolic Ca 2+ stimulate mitochondrial biogenesis in muscle cells. Am J Physiol Endocrinol Metab 283, E1040–E1045.

Paulsen G, Cumming KT, Holden G, Hallen J, Ronnestad BR, Sveen O, Skaug A, Paur I, Bastani NE, Ostgaard HN, Buer C, Midttun M, Freuchen F, Wiig H, Ulseth ET, Garthe I, Blomhoffi R, Benestad HB & Raastad T. (2014). Vitamin C and E supplementation hampers cellular adaptation to endurance training in humans: a double-blind, randomised, controlled trial. J Physiol 592, 1887–1901.

Peake JM, Della Gatta P, Suzuki K & Nieman DC. (2015). Cytokine expression and secretion by skeletal muscle cells: regulatory mechanisms and exercise effiects. Exerc Immunol Rev 21, 8–25.

Perez-Riverol Y, Bai J, Bandla C, Garcia-Seisdedos D, Hewapathirana S, Kamatchinathan S, Kundu DJ, Prakash A, Frericks-Zipper A, Eisenacher M, Walzer M, Wang S, Brazma A & Vizcaino JA. (2022). The PRIDE database resources in 2022: a hub for mass spectrometry-based proteomics evidences. Nucleic Acids Res 50, D543–D552.

Perry CGR, Lally J, Holloway GP, Heigenhauser GJF, Bonen A & Spriet LL. (2010). Repeated transient mRNA bursts precede increases in transcriptional and mitochondrial proteins during training in human skeletal muscle. J Physiol 588, 4795–4810.

Place N, Ivarsson N, Venckunas T, Neyroud D, Brazaitis M, Cheng AJ, Ochala J, Kamandulis S, Girard S, Volungevicius G, Pauzas H, Mekideche A, Kayser B, Martinez-Redondo V, Ruas JL, Bruton J, Truffiert A, Lanner JT, Skurvydas A & Westerblad H. (2015). Ryanodine receptor fragmentation and sarcoplasmic reticulum Ca^2+^ leak after one session of high-intensity interval exercise. Proc Natl Acad Sci U S A 112, 15492–15497.

Powers SK & Jackson MJ. (2008). Exercise-induced oxidative stress: cellular mechanisms and impact on muscle force production. Physiol Rev 88, 1243–1276.

Powers SK, Radak Z, Ji LL & Jackson M. (2024). Reactive oxygen species promote endurance exercise-induced adaptations in skeletal muscles. J Sport Health Sci 13, 780–792.

Raschke S & Eckel J. (2013). Adipo-myokines: two sides of the same coin--mediators of inflammation and mediators of exercise. Mediators Inflamm 2013, 320724.

Reisman EG, Botella J, Huang C, Schittenhelm RB, Stroud DA, Granata C, Chandrasiri OS, Ramm G, Oorschot V, Caruana NJ & Bishop DJ. (2024a). Fibre-specific mitochondrial protein abundance is linked to resting and post-training mitochondrial content in the muscle of men. Nat Commun 15, 7677.

Reisman EG, Caruana NJ & Bishop DJ. (2024b). Exercise training and changes in skeletal muscle mitochondrial proteins: from blots to “omics”. Crit Rev Biochem Mol Biol 59, 221–243.

Richards AJ, Watanabe D, Yamada T, Westerblad H & Cheng AJ. (2025). Task-dependent mechanisms underlying prolonged low-frequency force depression. Exerc Sport Sci Rev 53, 41–47.

Ristow M, Zarse K, Oberbach A, Klöting N, Birringer M, Kiehntopf M, Stumvoll M, Kahn CR & Blüher M. (2009). Antioxidants prevent health-promoting effiects of physical exercise in humans. Proc Natl Acad Sci U S A 106, 8665–8670.

Schlittler M, Neyroud D, Tanga C, Zanou N, Kamandulis S, Skurvydas A, Kayser B, Westerblad H, Place N & Andersson DC. (2019). Three weeks of sprint interval training improved high-intensity cycling performance and limited ryanodine receptor modifications in recreationally active human subjects. Eur J Appl Physiol 119, 1951–1958.

Sies H, Berndt C & Jones DP. (2017). Oxidative Stress. Annu Rev Biochem 86, 715–748.

Sies H & Cadenas E. (1985). Oxidative stress: damage to intact cells and organs. Philos Trans R Soc Lond B Biol Sci 311, 617–631.

Skelly LE, Gillen JB, Frankish BP, MacInnis MJ, Godkin FE, Tarnopolsky MA, Murphy RM & Gibala MJ. (2021). Human skeletal muscle fiber type-specific responses to sprint interval and moderate-intensity continuous exercise: acute and training-induced changes. J Appl Physiol 130, 1001–1014.

Sperlich B, Matzka M & Holmberg HC. (2023). The proportional distribution of training by elite endurance athletes at diffierent intensities during diffierent phases of the season. Front Sports Act Living 5, 1258585.

Strobel NA, Peake JM, Matsumoto A, Marsh SA, Coombes JS & Wadley GD. (2011). Antioxidant supplementation reduces skeletal muscle mitochondrial biogenesis. Med Sci Sports Exerc 43, 1017–1024.

Tan R, Nederveen JP, Gillen JB, Joanisse S, Parise G, Tarnopolsky MA & Gibala MJ. (2018). Skeletal muscle fiber-type-specific changes in markers of capillary and mitochondrial content after low-volume interval training in overweight women. Physiol Rep 6, e13597.

Tripp TR, Frankish BP, Lun V, Wiley JP, Shearer J, Murphy RM & MacInnis MJ. (2022). Time course and fibre type-dependent nature of calcium-handling protein responses to sprint interval exercise in human skeletal muscle. J Physiol 600, 2897–2917.

Tyanova S, Temu T & Cox J. (2016). The MaxQuant computational platform for mass spectrometry-based shotgun proteomics. Nat Protoc 11, 2301–2319.

Vøllestad NK & Blom PC. (1985). Effiect of varying exercise intensity on glycogen depletion in human muscle fibres. Acta Physiol Scand 125, 395–405.

Wright DC, Geiger PC, Han DH, Jones TE & Holloszy JO. (2007). Calcium induces increases in peroxisome proliferator-activated receptor gamma coactivator-1α and mitochondrial biogenesis by a pathway leading to p38 mitogen-activated protein kinase activation. J Biol Chem 282, 18793–18799.

Wyckelsma VL, Levinger I, Murphy RM, Petersen AC, Perry BD, Hedges CP, Anderson MJ & McKenna MJ. (2017). Intense interval training in healthy older adults increases skeletal muscle [^3^H]ouabain-binding site content and elevates Na^+^,K^+^-ATPase α2 isoform abundance in Type II fibers. Physiol Rep 5, e13219.

Wyckelsma VL, Venckunas T, Brazaitis M, Gastaldello S, Snieckus A, Eimantas N, Baranauskiene N, Subocius A, Skurvydas A, Paasuke M, Gapeyeva H, Kaasik P, Paasuke R, Jurimae J, Graf BA, Kayser B, Place N, Andersson DC, Kamandulis S & Westerblad H. (2020). Vitamin C and E treatment blunts sprint interval training-induced changes in inflammatory mediator-, calcium-, and mitochondria-related signaling in recreationally active elderly humans. Antioxidants (Basel) 9, 879.

Wyckelsma VL, Venckunas T, Houweling PJ, Schlittler M, Lauschke VM, Tiong CF, Wood HD, Ivarsson N, Paulauskas H, Eimantas N, Andersson DC, North KN, Brazaitis M & Westerblad H. (2021). Loss of alpha-actinin-3 during human evolution provides superior cold resilience and muscle heat generation. Am J Hum Genet 108, 446–457.

Yfanti C, Åkerström T, Nielsen S, Nielsen AR, Mounier R, Mortensen OH, Lykkesfeldt J, Rose AJ, Fischer CP & Pedersen BK. (2010). Antioxidant supplementation does not alter endurance training adaptation. Med Sci Sports Exerc 42, 1388–1395.

