## Supplemental Figures and Table for "Antioxidant supplementation blunts the proteome response to three weeks of sprint interval training preferentially in human type 2 muscle fibres"

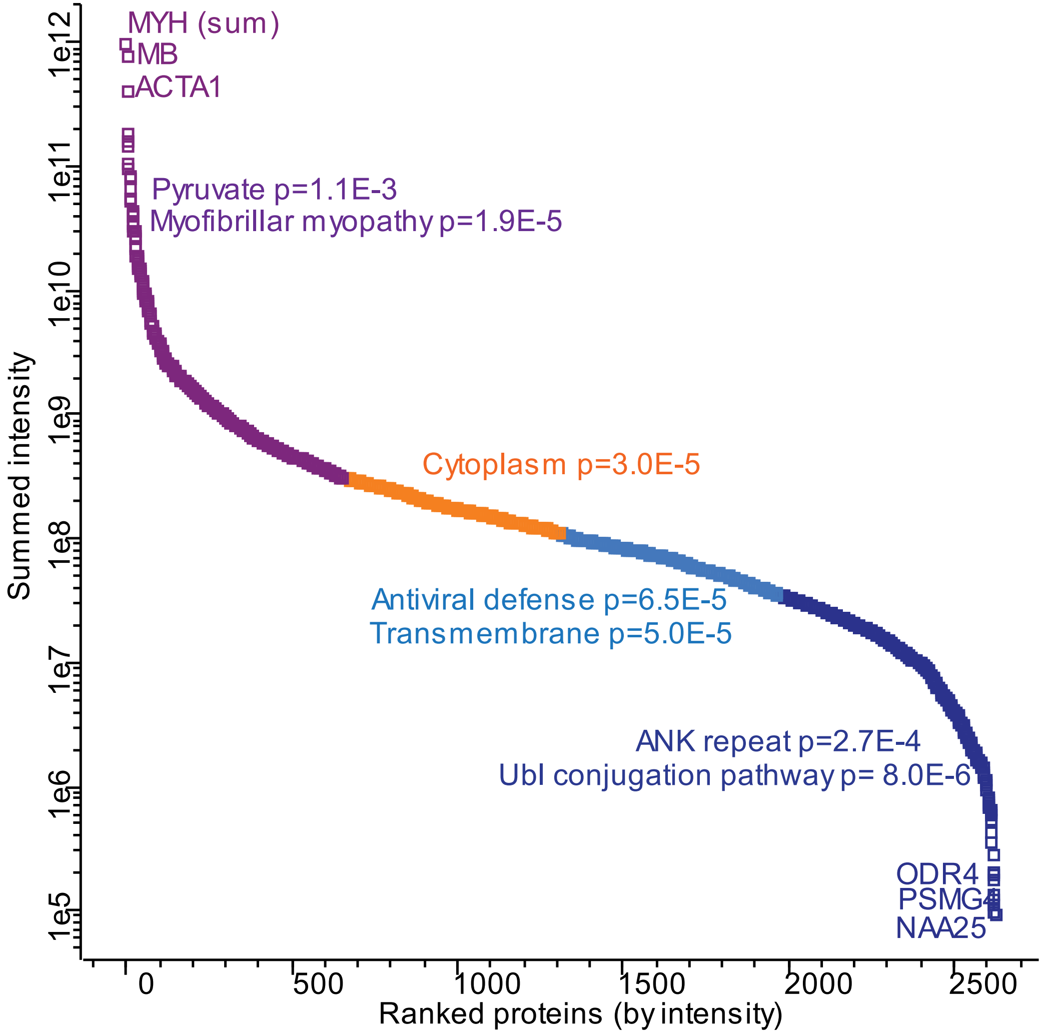

**Supplemental Figure 1**: Intensity distribution of the proteins quantified in the dataset, divided in quartiles (see different symbol color). The top two Uniprot keywords annotation enrichments in each quartile are shown next to the curve, with the corresponding p value (Fisher's exact test, FDR 0.02, Q2 only one enrichment). The gene name of the three most and least intense proteins is shown on top and bottom of the curve, respectively. Intensities of the adult MYH isoforms (slow: MYH7; fast: MYH2, MYH1, MYH4) were summed before ranking.

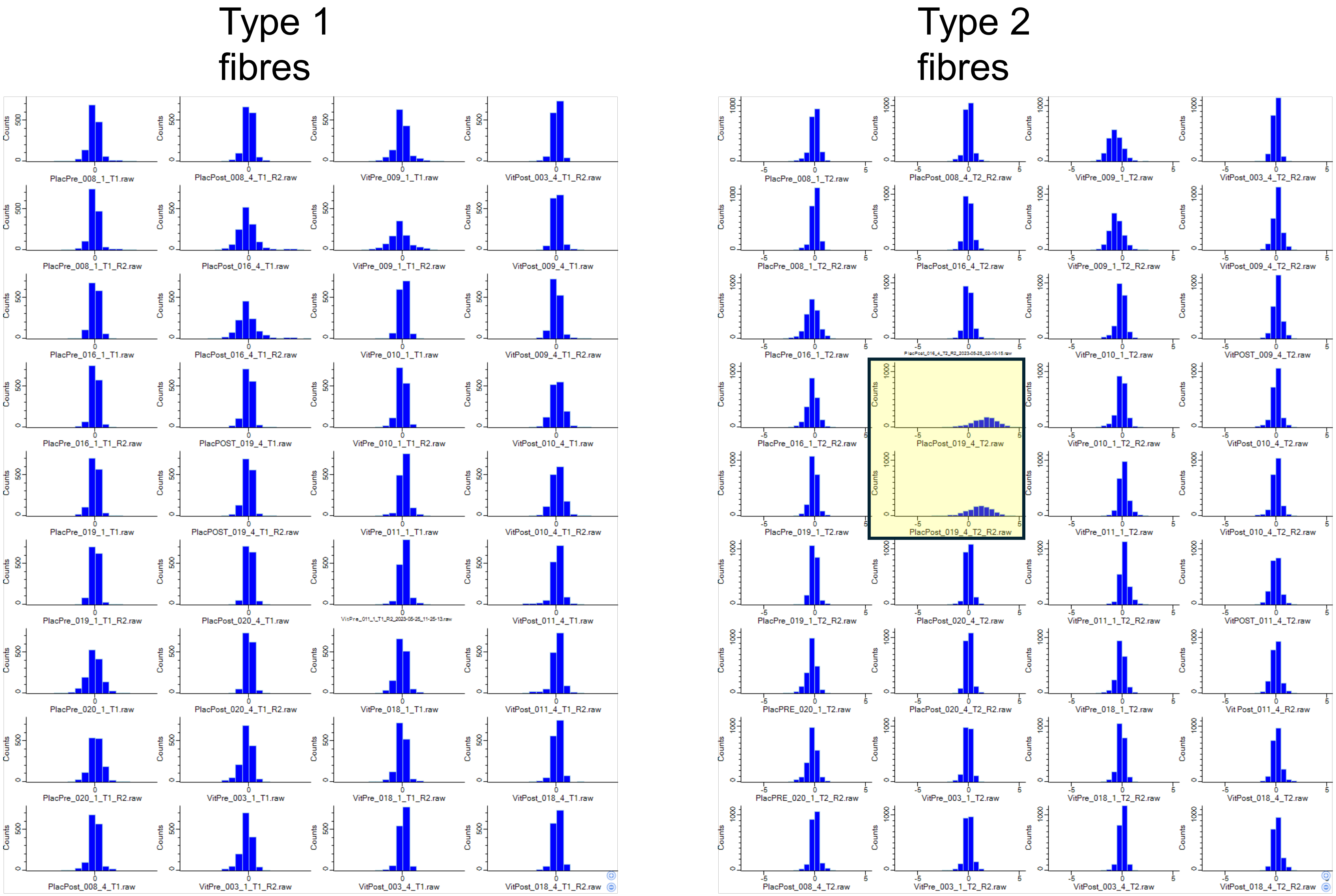

**Supplemental Figure 2**: Distribution histograms of $log_2$ transformed protein expression values normalized by subtracting mean values. Duplicate measurements were performed for each muscle sample. Samples from vitamin (Vit) and placebo (Pla) treated participants obtained before (Pre) and after (Post) the three weeks of SIT. Histograms show the expected clear peak and close to normal distribution for each sample with one exception: type 2 fibres PlaPost_019 (shaded box), hence the type 2 fibre data from this participant were excluded from further analyses.

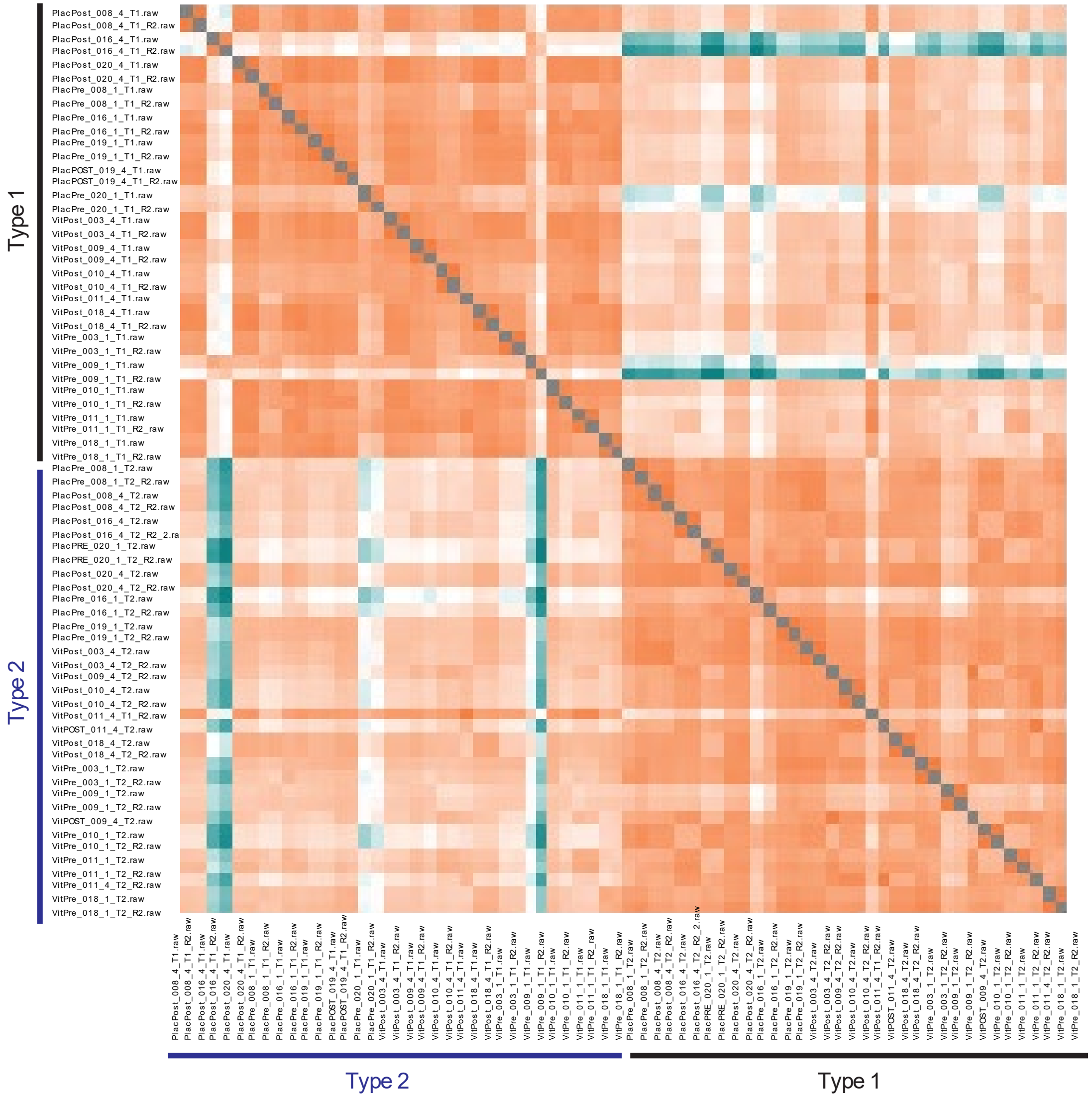

**Supplemental Figure 3**: Matrix of Pearson correlations between measured protein intensities of individual fibre pools in the dataset. Technical duplicates of each muscle fibre pool are shown as individual runs

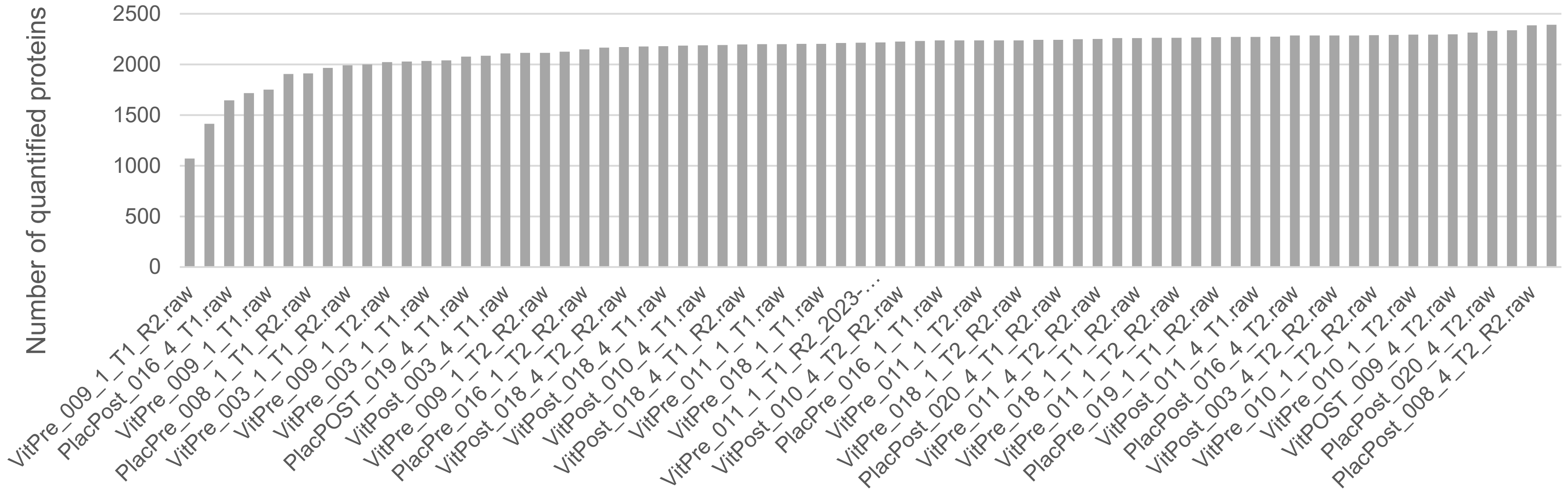

**Supplemental Figure 4**: Number of quantified proteins in individual MS runs for each fiber pool sample. File names correspond to those of the raw files.

**Supplemental Table 1. Mitochondrial proteins selected with KEGG name filtering using the terms “oxidative phosphorylation” and “citrate cycle”, and excluding proteins that according to MitoCarta 3.0 are not directly involved in aerobic ATP production or not not localized in mitochondria.**

| | **Gene** | **Protein description** |
|---|---|---|
| **Pyruvate and tricarboxylic acid cycle** | ACO2 | Aconitate hydratase, mitochondrial |
| | CS | Citrate synthase, mitochondrial |
| | DLAT | Dihydrolipoyllysine-residue acetyltransferase component of pyruvate dehydrogenase complex, mitoch. |
| | DLD | Dihydrolipoyl dehydrogenase, mitochondrial |
| | DLST | Dihydrolipoyllysine-residue succinyltransferase comp. of 2-oxoglutarate dehydrogenase compl, mitoch |
| | FH | Isoform of P07954, Isoform Cytoplasmic of Fumarate hydratase, mitochondrial |
| | IDH2 | Isocitrate dehydrogenase [NADP], mitochondrial |
| | IDH3A | Isocitrate dehydrogenase [NAD] subunit alpha, mitochondrial |
| | IDH3B | Isoform of O43837, Isoform A of Isocitrate dehydrogenase [NAD] subunit beta, mitochondrial |
| | IDH3G | Isocitrate dehydrogenase [NAD] subunit gamma, mitochondrial |
| | MDH2 | Malate dehydrogenase, mitochondrial |
| | OGDH | 2-oxoglutarate dehydrogenase, mitochondrial |
| | PDHA1 | Pyruvate dehydrogenase E1 component subunit alpha, somatic form, mitochondrial |
| | PDHB | Pyruvate dehydrogenase E1 component subunit beta, mitochondrial |
| | PDHX | Pyruvate dehydrogenase protein X component, mitochondrial |
| | SUCLA2 | Succinate--CoA ligase [ADP-forming] subunit beta, mitochondrial |
| | SUCLG1 | Succinate--CoA ligase [ADP/GDP-forming] subunit alpha, mitochondrial |
| | SUCLG2 | Succinate--CoA ligase [GDP-forming] subunit beta, mitochondrial |
| **Complex 1** | MT-ND1 | NADH-ubiquinone oxidoreductase chain 1 |
| | MT-ND2 | NADH-ubiquinone oxidoreductase chain 2 |
| | MT-ND4 | NADH-ubiquinone oxidoreductase chain 4 |
| | MT-ND5 | NADH-ubiquinone oxidoreductase chain 5 |
| | NDUFA1 | NADH dehydrogenase [ubiquinone] 1 alpha subcomplex subunit 1 |
| | NDUFA10 | NADH dehydrogenase [ubiquinone] 1 alpha subcomplex subunit 10, mitochondrial |
| | NDUFA11 | NADH dehydrogenase [ubiquinone] 1 alpha subcomplex subunit 11 |

| | | |
|---|---|---|
| **Complex 1, cont.** | NDUFA12 | NADH dehydrogenase [ubiquinone] 1 alpha subcomplex subunit 12 |
| | NDUFA13 | NADH dehydrogenase [ubiquinone] 1 alpha subcomplex subunit 13 |
| | NDUFA2 | NADH dehydrogenase [ubiquinone] 1 alpha subcomplex subunit 2 |
| | NDUFA3[1] | NADH dehydrogenase [ubiquinone] 1 alpha subcomplex subunit 3 |
| | NDUFA5 | NADH dehydrogenase [ubiquinone] 1 alpha subcomplex subunit 5 |
| | NDUFA6 | NADH dehydrogenase [ubiquinone] 1 alpha subcomplex subunit 6 |
| | NDUFA7 | NADH dehydrogenase [ubiquinone] 1 alpha subcomplex subunit 7 |
| | NDUFA8 | NADH dehydrogenase [ubiquinone] 1 alpha subcomplex subunit 8 |
| | NDUFA9 | NADH dehydrogenase [ubiquinone] 1 alpha subcomplex subunit 9, mitochondrial |
| | NDUFAB1 | Acyl carrier protein, mitochondrial |
| | NDUFB1 | NADH dehydrogenase [ubiquinone] 1 beta subcomplex subunit 1 |
| | NDUFB10 | NADH dehydrogenase [ubiquinone] 1 beta subcomplex subunit 10 |
| | NDUFB11 | NADH dehydrogenase [ubiquinone] 1 beta subcomplex subunit 11, mitochondrial |
| | NDUFB3 | NADH dehydrogenase [ubiquinone] 1 beta subcomplex subunit 3 |
| | NDUFB4 | NADH dehydrogenase [ubiquinone] 1 beta subcomplex subunit 4 |
| | NDUFB5 | NADH dehydrogenase [ubiquinone] 1 beta subcomplex subunit 5, mitochondrial |
| | NDUFB6[1] | NADH dehydrogenase [ubiquinone] 1 beta subcomplex subunit 6 |
| | NDUFB7 | NADH dehydrogenase [ubiquinone] 1 beta subcomplex subunit 7 |
| | NDUFB8 | NADH dehydrogenase [ubiquinone] 1 beta subcomplex subunit 8, mitochondrial |
| | NDUFB9 | NADH dehydrogenase [ubiquinone] 1 beta subcomplex subunit 9 |
| | NDUFC1[1] | NADH dehydrogenase [ubiquinone] 1 subunit C1, mitochondrial |
| | NDUFC2 | NADH dehydrogenase [ubiquinone] 1 subunit C2 |
| | NDUFS1 | NADH-ubiquinone oxidoreductase 75 kDa subunit, mitochondrial |
| | NDUFS2 | NADH dehydrogenase [ubiquinone] iron-sulfur protein 2, mitochondrial |
| | NDUFS3 | NADH dehydrogenase [ubiquinone] iron-sulfur protein 3, mitochondrial |
| | NDUFS4 | NADH dehydrogenase [ubiquinone] iron-sulfur protein 4, mitochondrial |
| | NDUFS5 | NADH dehydrogenase [ubiquinone] iron-sulfur protein 5 |
| | NDUFS6 | NADH dehydrogenase [ubiquinone] iron-sulfur protein 6, mitochondrial |
| | NDUFS7 | Isoform of O75251, Isoform 2 of NADH dehydrogenase [ubiquinone] iron-sulfur protein 7, mitochondrial |
| | NDUFS8 | NADH dehydrogenase [ubiquinone] iron-sulfur protein 8, mitochondrial |

| | | |
|---|---|---|
| **Complex 1, cont.** | NDUFV1 | NADH dehydrogenase [ubiquinone] flavoprotein 1, mitochondrial |
| | NDUFV2 | NADH dehydrogenase [ubiquinone] flavoprotein 2, mitochondrial |
| | NDUFV3 | NADH dehydrogenase [ubiquinone] flavoprotein 3, mitochondrial |
| | NDUFV3-2 | Isoform of P56181, Isoform 2 of NADH dehydrogenase [ubiquinone] flavoprotein 3, mitochondrial |
| **Complex 2** | SDHA | Succinate dehydrogenase [ubiquinone] flavoprotein subunit, mitochondrial |
| | SDHB | Succinate dehydrogenase [ubiquinone] iron-sulfur subunit, mitochondrial |
| | SDHC | Succinate dehydrogenase cytochrome b560 subunit, mitochondrial |
| | SDHD | Succinate dehydrogenase [ubiquinone] cytochrome b small subunit, mitochondrial |
| **Complex 3** | CYC1 | Cytochrome c1, heme protein, mitochondrial |
| | MT-CYB | Cytochrome b |
| | UQCR10 | Cytochrome b-c1 complex subunit 9 |
| | UQCRB | Cytochrome b-c1 complex subunit 7 |
| | UQCRC1 | Cytochrome b-c1 complex subunit 1, mitochondrial |
| | UQCRC2 | Cytochrome b-c1 complex subunit 2, mitochondrial |
| | UQCRFS1 | Cytochrome b-c1 complex subunit Rieske, mitochondrial |
| | UQCRH | Cytochrome b-c1 complex subunit 6, mitochondrial |
| | UQCRQ | Cytochrome b-c1 complex subunit 8 |
| **Complex 4** | COX11[2] | Cytochrome c oxidase assembly protein COX11, mitochondrial |
| | COX17 | Cytochrome c oxidase copper chaperone |
| | COX4I1 | Cytochrome c oxidase subunit 4 isoform 1, mitochondrial |
| | COX5A | Cytochrome c oxidase subunit 5A, mitochondrial |
| | COX5B | Cytochrome c oxidase subunit 5B, mitochondrial |
| | COX6A2 | Cytochrome c oxidase subunit 6A2, mitochondrial |
| | COX6B1 | Cytochrome c oxidase subunit 6B1 |
| | COX6C | Cytochrome c oxidase subunit 6C |
| | COX7A1 | Cytochrome c oxidase subunit 7A1, mitochondrial |
| | COX7A2 | Cytochrome c oxidase subunit 7A2, mitochondrial |

| | | |
|---|---|---|
| | MT-CO1 | Cytochrome c oxidase subunit 1 |
| | MT-CO2 | Cytochrome c oxidase subunit 2 |
| **Complex 4, cont.** | MT-CO3 | Cytochrome c oxidase subunit 3 |
| | NDUFA4 | Cytochrome c oxidase subunit NDUFA4 |
| **Complex 5** | ATP5F1A | ATP synthase subunit alpha, mitochondrial |
| | ATP5F1B | ATP synthase subunit beta, mitochondrial |
| | ATP5F1C | ATP synthase subunit gamma, mitochondrial |
| | ATP5F1D | ATP synthase subunit delta, mitochondrial |
| | ATP5F1E | ATP synthase subunit epsilon, mitochondrial |
| | ATP5ME | ATP synthase subunit e, mitochondrial |
| | ATP5MF | ATP synthase subunit f, mitochondrial |
| | ATP5MG | ATP synthase subunit g, mitochondrial |
| | ATP5PB | ATP synthase F(0) complex subunit B1, mitochondrial |
| | ATP5PD | ATP synthase subunit d, mitochondrial |
| | ATP5PF | ATP synthase-coupling factor 6, mitochondrial |
| | ATP5PO | ATP synthase subunit O, mitochondrial |
| | MT-ATP6 | ATP synthase subunit a |

[1]Only detected in type 1 fibres; [2]only detected in type 2 fibres
